# Harnessing droplet microfluidics and morphology-based deep learning for the label-free study of polymicrobial-phage interactions

**DOI:** 10.1101/2025.03.24.644924

**Authors:** Anuj Tiwari, An Mei Daniels, Robyn Manley, Fabrice Gielen

## Abstract

Evaluating the impact of bacteriophages on bacterial communities is required to assess the future utility of phage therapy. Methods able to study bacterial polycultures in the presence of phages are useful to mimic evolutionary pressures found in natural environments and recapitulate complex ecological contexts. Bacteriophages can drive rapid genetic and phenotypic changes in host cells. However, the presence of other bacteria can also impact bacterial densities and community structure, and classical methods remain lengthy and resource intensive. Here, we introduce a microdroplet-based encapsulation method in which bacterial co-cultures are imaged using Z-stack brightfield microscopy. The method relies on automated droplet imaging using a novel AI-based autofocus function, coupled with morphology-based deep learning models for accurate identification of two morphologically distinct bacterial species. We show that we can monitor the relative growth dynamics of *P. aeruginosa* and *S. aureus* growing in 11 picolitre droplets for up to 24 hours. We demonstrate quantification of growth rates, bacterial densities and lysis dynamics of the two species without the need for plating. We show that a potent lytic phage of *P. aeruginosa* can either fully lyse the initial *P. aeruginosa* population or keep its density low long-term when in the presence of *S. aureus*.

## Introduction

Bacteriophages are abundant in nature and major drivers of bacterial evolution. The use of phages to lyse specific bacteria has recently seen a resurgence in interest as a promising approach for tackling antibiotics resistance and diseases beyond infections. In addition to potential live drugs for human treatment, phages can find applications in decontamination of crops, animal feeds or hospital settings. However, bacterial phenotypes triggered by phage exposure in monocultures can significantly differ from those found in polymicrobial communities^1^. For example, rapid resistance acquisition and high fitness of resistant bacteria have often been reported when using monocultures. Yet, it is known that this evolutionary outcome of bacteria-phage interactions can be fundamentally altered by the presence of a polymicrobial community and this could be a key consideration in the development of future phage-based therapies^2^. It is increasingly clear that resistance mechanisms are context dependent and that phage resistance arise differently *in vitro* and in natural environments. For example, surface modifications and receptor shedding have been reported in laboratory conditions while CRISPR-Cas based defense systems are seen more often in polymicrobial communities due to fitness costs of certain types of resistance^3^. Beyond how resistance eventually arise, the question is whether microbial diversity promote or impede phage infections and how polymicrobial-phage interactions shape community structure.

In a study by Testa *et al.*, a sensitive *P. aeruginosa* strain PAO1 was cultured with its lytic phage in the presence of an insensitive *P. aeruginosa* strain PA14. It was found that inter-strain competition reduced resistance evolution in the susceptible strain ^4^. In another study by Harcombe *et al.* with *E. coli B* and *Salmonella enterica*, T5 and T7 phages kept *E. coli* density levels low in the two-species cultures although densities returned to the same levels as without phages in *E. coli* monocultures^5^. It was recently shown that phages are most likely to affect community composition when encounter rates with susceptible hosts is high, over short timescales, and in relatively simple communities^6^.

We chose to co-culture of *P. aeruginosa* (PA 14 ΔflgK, referred to as PA) and *S. aureus* (MSSA476, referred to as SA) as a model two-species community. PA is a common cause of nosocomial infections, accounting for 10–20% of infections in most hospitals and the combination with SA has high relevance to cystic fibrosis lung environments and chronic wound infections ^7^. Many reports found that PA strongly outcompetes SA in liquid and biofilm cultures ^8,9^. This is in part thanks to the secretion of several virulence and quorum sensing compounds including pyocyanins and polyphosphates which disrupts *S. aureus* respiratory processes and can cause oxidative stress^10^. However, other reports found culturing media conditions that support *P. aeruginosa* and *S. aureus* coexistence *in vitro*^11^*. S aureus* growth within biofilms has also been shown to display high dependency on oxygen availability^12^.

Current methods for studying polymicrobial cultures are lengthy and labour intensive, typically based on liquid cultures followed by quantification of the relative abundance of either bacterial species found by plating on suitable selective agar plates. Standard optical density measurements cannot report on the relative growth of multiple species in the same mixed culture ^13^. Other methods include flow cytometry of fluorescently labelled cells, or molecular techniques such as targeted sequencing, multiplex qPCR-based methods^14^. Imaging methods can be used to identify bacteria and detect specific phenotypes^15^. However, most of the current imaging methods require cells to grow in single layers with limited relevance to 3D bacterial niches^16,17^. Counting and identifying motile cells swimming in 3D environments is more challenging due to the rapid spreading and random orientation of bacteria at any time. Microfluidic encapsulation methods enable facile trapping of individual cells necessary for long-term imaging^18^. The use of microfluidic methods enables rapid generation of co-cultures with precise control over cellular encapsulation parameters (e.g. volumes, trapping locations) and the creation of diverse microbial communities with starting bacteria numbers following Poisson statistics ^19^. Microbial interactions have been studied in droplets, for instance with the co-encapsulation of symbiotic fluorescent bacteria^20^. These studies have required fluorescent reporters and relied on quantifying overall fluorescent levels to estimate bacterial densities over time ^21,22^.

We have previously demonstrated label-free counting of *E. coli* cells trapped in non-contacting static microdroplets with a low-throughput approach that enabled tracking of individual bacterial division events ^23^. Here we expand the method to two-species cultures using rod-shaped (PA) and cocci-shaped (SA) cells trapped within shallow water-in-oil microdroplets in which strains interact. We show that deep learning object detectors can accurately count the number of cells of both species based on their morphological signatures ^24^. A Z-stacking imaging method enables accurate counting across the 3D bacterial niches. We further introduce an AI-based autofocus function enabling stable and reproducible focus over timescales of hours while the microscope stage and objective undergo repetitive XY and Z motion respectively. Previous studies have investigated the use of autofocus methods using convolutional neural networks to determine the focal plane of an image in a single shot, and identify the distance and direction needed to return to focus^25^. This is more efficient than other implementations using multiple slices a known distance apart to extrapolate focal distance^26^. However, all of the networks were trained on stained or auto-fluorescent tissue samples, where all cells were in the same plane^27–29^. Our autofocus system was trained on 5 separate classes corresponding to 5 different focal range for the droplets and a feedback loop was implemented to converge towards best focus for every droplet screened. This enabled us to collect reproducible experimental data with non-fluorescent bacterial cells for long-term time-lapse imaging.

Using the platform, we monitored the relative growth dynamics of *P. aeruginosa* and *S. aureus for* up to 20 co-cultures per experiment growing in confined 12 picolitre droplets for up to 24 hours. We challenged monocultures and polycultures to a lytic phage of *P. aeruginosa* and demonstrate quantification of growth rates, relative bacterial densities and lysis dynamics for the two species.

Our data show that in the absence of phage, SA populations in a mixed SA+PA community had usually, but not always, reduced growth rates compared to monoculture controls. The presence of PA phage P278 fully lysed or suppressed PA while promoting *S. aureus* growth in two-species cultures over timescales of over 5 hours suggesting long-term growth suppression effect.

## 1. Methods

### 1.1 Bacterial strain and phage lysate preparation

*Pseudomonas aeruginosa* strain *PA14-ΔflgK* (gram negative) and *Staphylococcus aureus* strain *MSSA-476* (gram positive) were chosen as the model strains for this polymicrobial study. Lysogeny Broth (LB) agar plates with single colonies of each strain were obtained from glycerol stocks stored at −80°C. To initiate an experiment, a single colony from each plate was picked and added to a sterile culture tube containing 5 mL of LB media (10 g/L tryptone, 5 g/L yeast extract, 10 g/L NaCl) and incubated overnight at 37°C with shaking at 200 rpm. Different volumes of overnight culture were then added to 5 mL of fresh LB media for further cultivation until a desired OD_600_ (optical density measured at 600 nm) was obtained depending on specific experimental requirements.

The number of cells in droplets follows Poisson distribution. This allows us to predict the average number of cells in each droplet system based on OD_600_ values. A broad range of experiments were conducted to obtain data on individual strains as well as their combination starting at different OD_600_ values to study strain fitness. Desired OD_600_ values for growth experiments were chosen between 0.100-0.400 to encapsulate fewer than 5 cells in each droplet. Lysis experiments were conducted at OD_600_ values between 0.6-0.8 to ensure that each droplet had at least 10 cells. Table 1 shows all experimental conditions for growth experiments. Standard colony forming unit (CFU) assays were also carried out to obtain specific population numbers pertaining to the OD_600_ values.

**Table 1:** Growth experiment conditions for individual and polymicrobial droplets.

| Experiment | OD<sub>600</sub> PA14 | OD<sub>600</sub> MSSA476 |
| --- | --- | --- |
| PA14 Growth 1 | 0.11 | - |
| PA14 Growth 2 | 0.14 | - |
| PA14 Growth 3 | 0.14 | - |
| MSSA 476 Growth 1 | - | 0.14 |
| MSSA 476 Growth 2 | - | 0.13 |
| MSSA 476 Growth 3 | - | 0.14 |
| Polymicrobial Growth 1 | 0.10 | 0.16 |
| Polymicrobial Growth 2 | 0.31 | 0.40 |

Phage lysate P278 was preserved in SM (Storage and Maintenance) buffer (0.1 M NaCl, 8 mM MgSO4·7H2O, 50 mM Tris-Cl, pH 7.5, 0.01% gelatin) at 4°C, with phage titer of 8 x 10^9^ PFU/mL (PFU, plaque-forming unit). For single strain as well as polymicrobial lysis experiments, different volumes of strains cultured in exponential phase were mixed with the cell lysate to obtain desirable multiplicity of infection (MOI). MOI is defined as the ratio between the number of phage particles and number of bacterial cells.

### 1.2 Microfluidic chip fabrication

The two-layer microfluidic device was produced using soft lithography and a high-resolution acetate mask (Microlithography Services). The procedure involved patterning negative photoresist SU8 TF6000 (MicroChem) on a silicon wafer, which was exposed to UV light through a transparent film mask. The device comprises two layers: the first layer includes 3 μm-high pillars, while the second layer contains circular microdroplet traps, also 3 μm in height, with a diameter of 60 μm. Further details on the procedure can be found in a previous publication ^23^. A mixture of PDMS (polydimethylsiloxane, Ellsworth) and a curing agent at a 10:1 ratio was poured onto the patterned wafer after being degassed. To prevent droplet evaporation during time-lapse experiments, a small coverslip (0.15 mm thick) was placed over the trap array before PDMS curing. The wafer was cured at 70°C for 120 minutes. After curing, the PDMS was cut out and 1 mm inlet holes were punched (Kai Medical) for oil and bacterial culture media. Plasma treatment (Diener Zepto) was used to bond the PDMS to a thin coverslip (22 × 50 mm, 0.13–0.17 mm thick). Finally, a 1% (v/v) solution of trichloro(1H,1H,2H,2H-perfluorooctyl) silane (Merck) in HFE-7500 oil was flushed through the device, followed by incubation at 70°C for a minimum of 30 minutes.

### 1.3 Anchored droplet generation and trapping

The microfluidic device was positioned on the microscope stage and secured with scotch tape. Droplets were formed on-chip through phase change, where the aqueous phase containing bacterial cells or a bacteria-phage mixture was replaced by the oil phase, creating anchored droplets at the designed traps. The continuous phase was composed of 1% (w/v) 008-Fluorosurfactant (RAN Biotechnologies) in HFE-7500 oil (Fluorochem). The aqueous phase included either bacterial strains in LB broth for growth experiments or bacterial strains in LB broth with P278 phages for lysis-based experiments. A schematic of the microscope setup is shown in Figure 1. To initiate the experiments, the aqueous and oil phases were loaded into PTFE tubing (SLS) connected to 1 mL plastic syringes (BD Plastipak), which were mounted on syringe pumps (Nemesys, Cetoni). A volume between 100-200 μL of bacterial culture or bacteria-phage mix was aspirated into one syringe for injection into the chip. The device was first primed with oil to expel any air from the trapping chambers. Once the oil flow was stopped, the bacterial or bacteria-phage solution was introduced until the entire trapping array was filled (Figure 1B). The flow of the bacterial solution was then halted, and oil was reintroduced to flush the cell sample, creating droplets of the immobilized cell sample in the circular traps, as shown in Figure 1B.

**Figure 1:**
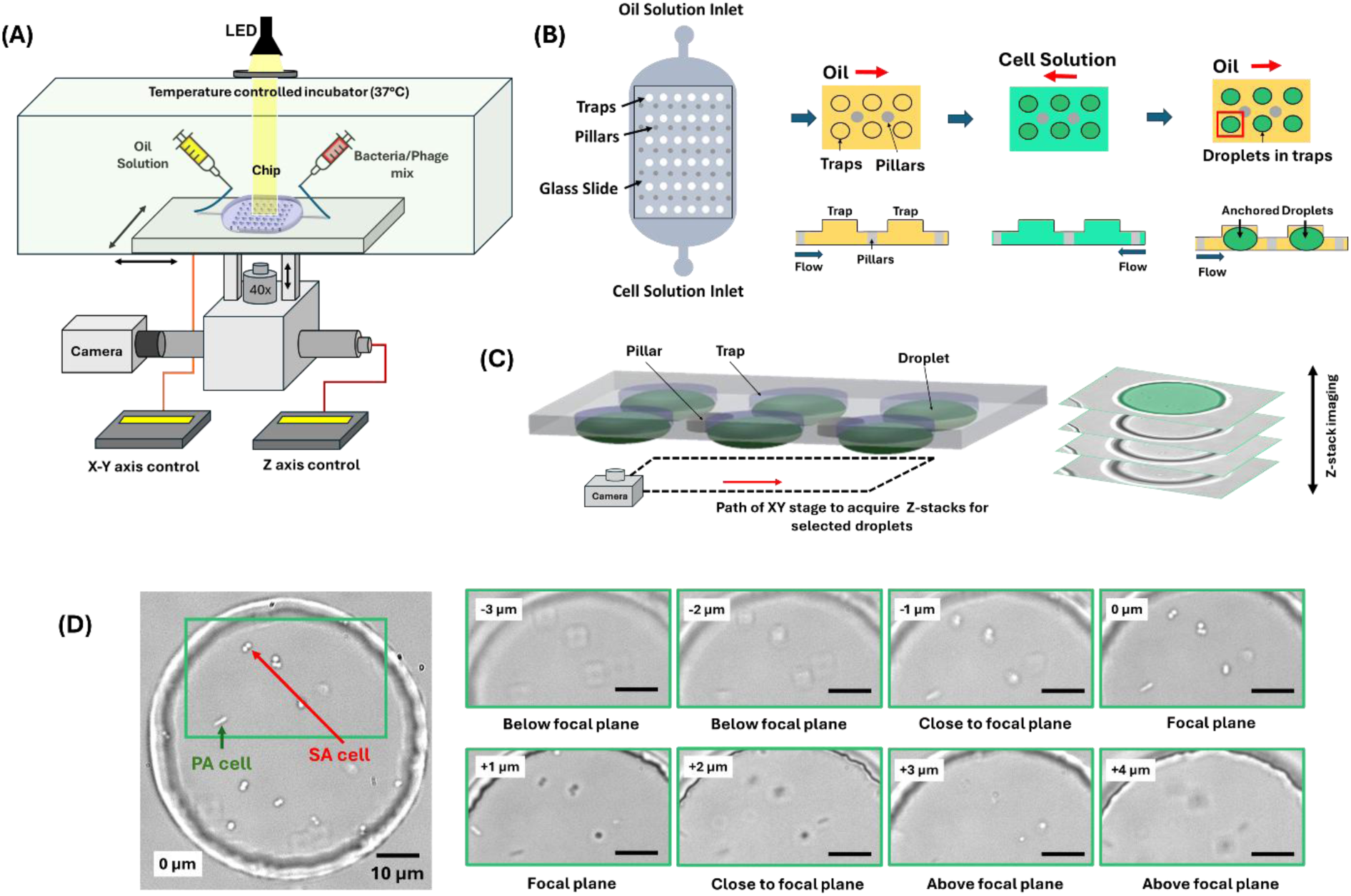
Microfluidics and imaging setup to study bacterial growth and bacteriophage interactions in anchored droplets (A) Schematic of the microfluidic experimental setup, including an inverted microscope equipped with a 40x magnification lens, a temperature-controlled incubator (37°C), a microfluidic chip comprising oil and bacterial/phage solution inputs, and an integrated camera (B) Illustration of the droplet generation process within the microfluidic chip, highlighting key stages of droplet formation using multiple flow directions, resulting in an anchored droplet array. (C) Automated imaging of multiple droplets using XY stage motion and Z-stacking to acquire imaging data to generate accurate counts (D) High-resolution microscopy images of bacterial cells (PA and SA) encapsulated in droplets, with scale bars of 10 µm. The green box highlights an example region of interest. Sequential microscopy images of a droplet loaded with bacteria, taken at different Z-axis positions (from −3 µm to +4 µm), showcasing the 3D positioning and morphological features at different focal planes for each species of bacterial cells.

**Figure 2:**
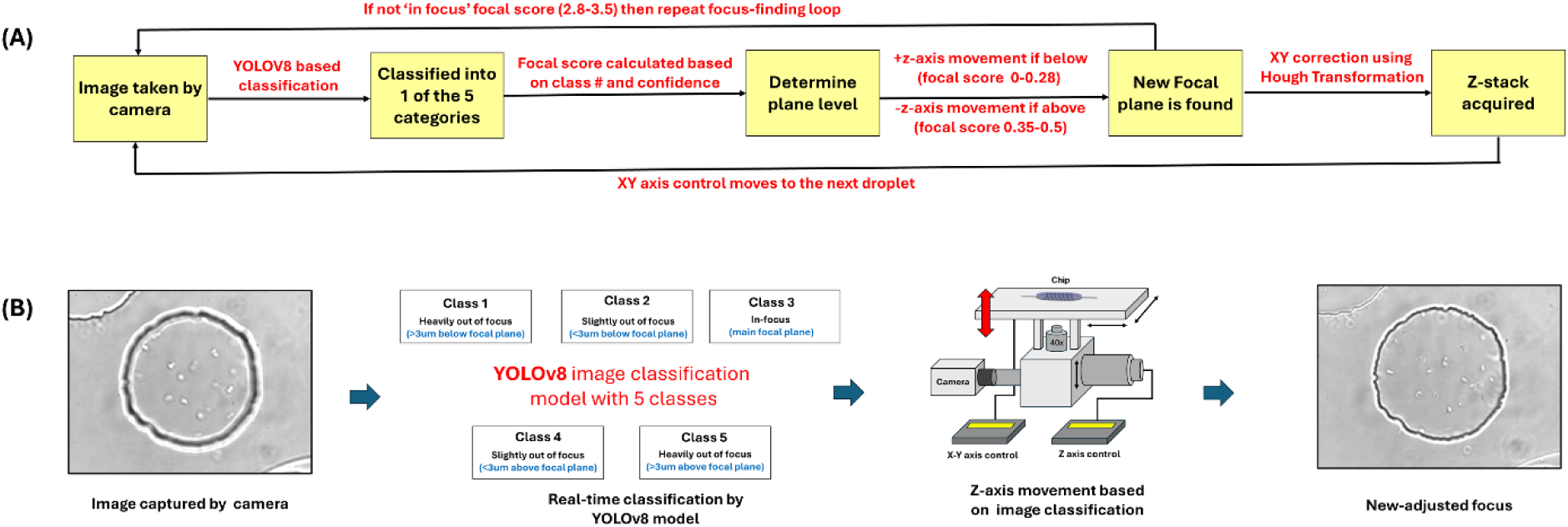
Autofocus using YOLOv8-based deep learning. (A) The workflow illustrates an automated focal adjustment system for droplet imaging using a YOLOv8-based classification model. Images are captured at 40x magnification and classified into five categories: heavily out of focus (>3 µm below or above the focal plane), slightly out of focus (<3 µm below or above the focal plane), and in-focus (close to the main focal plane). A focal score is calculated based on the classification and confidence level (see equation 1). An iterative focus adjustment loop is implemented if the score is not within the in-focus range (corresponding to a score of 2.8–3.5, with a score of 3 being the predicted best focus). The Z-axis is adjusted accordingly, with positive movement for below-focus scores (0–0.28) and negative movement for above-focus scores (0.35–0.5), until the correct focal plane is located. Subsequently, droplet centering in the XY plane is performed using Hough Transformation to identify circular traps, and a Z-stack of the droplet was acquired. (B) The bottom images illustrate the classification process, transitioning from an out-of-focus to an in-focus image. Movie S1 shows the functioning of the focus correction.

### 1.4 Imaging and focus correction using deep learning

The droplet imaging system developed in this study integrates automated time-lapse image acquisition, focal plane correction using deep learning, and axial drift (XY axis) compensation using Hough transform analyses. This system enables the precise imaging of individual droplets in a dynamic environment (i.e. with the droplet chip mounted on a moveable platform), ensuring that both the focal plane and droplet imaging conditions are maintained throughout long-duration experiments.

An inverted Olympus IX73 microscope with a motorized XY stage was used to image the bacterial cells trapped in different chambers on chip. The microscope was equipped with a white LED (CoolLED pE-100) and featured a bolt-on motorized Z-focus drive (PS3H122R, Prior Scientific) controlled by a ProScan III focus controller (Prior Scientific). Imaging was conducted using a 40x objective with a numerical aperture of 0.45 (UPLFLN40X-2, Olympus). The microscope was installed on a vibration-damping platform (Newport VIP320X1218-50140). A monochrome industrial USB camera (DMK 37AUX287, The Imaging Source) was used to capture images of the droplets at a resolution of 640 x 480 pixels, with 8 bits encoding. X and Y coordinates of selected droplets to be imaged within the droplet array were stored in a text file using a python script. Using another custom python script, the camera captured Z-slices spanning from above to below the best focal plane to generate a 3D stack for each droplet. The motorized stage and motorized Z-focus were used for precise X, Y, and Z movements during the imaging process with customizable focal adjustment steps.

To ensure consistent Z-stacks across all droplets, a pre-trained YOLOv8 deep learning-based image classification model was used to analyze initial droplet images and determine the current focal level. The model was specifically trained to classify focus levels into five categories (AboveGreater3, AboveLess3, InFocus, BelowGreater3, BelowLess3). The use of two different classes for slightly out-of-focus (<3µm) and greatly out-of-focus (>3µm) both above and below the plane allowed for greater accuracy in detecting location in the Z-stack, as droplets slightly out of focus in either direction were visually more similar to each other than very out-of-focus drops, regardless of direction. The categories were then remapped to an integer between 1 and 5 based on their respective classes for focus adjustment purposes. The system then computed a score weighted by confidence values based on model outputs to determine how far the image was from the best focus. We used the following equation to compute the score:

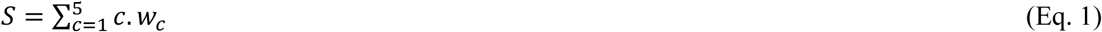

Where w_c_ is the confidence for each class c. If the score belonged to a predefined range of 2.8 ≤ *S* ≤ 3.5, the image was considered ‘in-focus’), as found empirically. Otherwise, the stage was adjusted by a pre-defined step size (typically 0.8 micron), and a newly acquired image was used to calculate an updated S score until optimal focus was achieved. The objective was moved upwards by 0.8 µm for scores below 2.8 and 0.8 µm upwards for values above 3.5.

The system recorded a history of focus movements over time (Figure S1). If the focus drifted beyond a pre-defined maximum number of iterations (corresponding to approx. 4 microns) for an extended period, the system detected this as focus instability and reversed the focus direction to attempt refocusing. Once focus was attained, a Hough transform was employed to detect the circular droplet trap and correct any axial drift of the droplet’s center position. The system first captured an image of the droplet and applied the Hough transform to identify circular patterns within the image. The detected center of the trap was compared to the expected initial position, and the system computed the necessary x and y adjustments to align the microscope stage. The stage was then moved to these updated coordinates, ensuring accurate droplet positioning over time (see Movie S1). The Python script also allowed modification of number of time points, the duration between each time point as well as number of images and spacing between slices in the Z-stack for each time point. This method enabled robust, automated droplet imaging with real-time corrections for focus drift and axial misalignment, ensuring high-quality imaging data over long experimental durations of over 12 hours.

### 1.5 Deep learning model generation for cell morphology detection

YOLOv5 (You Only Look Once, Version 5) is a single stage object detection model based on convolutional neural networks (CNN) developed by Ultralytics ^30^. It is widely used for detecting objects of interest in an image or a video at high speed and accuracy. YOLOv5 was chosen for object detection-based applications for this study as it outperforms many versions of object detection models such as YOLOv4, Mask R-CNN, R-CNN, RetinaNET and Single Shot MultiBox Detector ^31^.

YOLOv5x (extra-large) was chosen as the base model as it provides the highest detection accuracy compared to other models in the YOLOv5 family. In this study, we trained 3 different models of YOLOv5x for detecting bacterial strains with different morphologies (see Table 3). Some of the models were trained using transfer learning. Transfer learning is a machine learning technique where a model developed for one task is reused as the starting point for a model on a different, but related, task. The models were trained on Google COLAB webserver and the training was completed within a few minutes. All training parameters are reported in the supplementary section S1 with mAP, training and validation specifications.

**Table 2:** Lysis experiment conditions for individual and polymicrobial droplets.

| Experiment | CFU Count<br>(PA14)/ml | OD<sub>600</sub> |  | Phage titer<br>concentration<br>(Phage - P278)<br>(PFU/ml) | MOI |
| --- | --- | --- | --- | --- | --- |
|  |  | PA14 | MSSA476 |  |  |
| PA14 lysis 1 | 4.00 x 10<sup>8</sup> | 0.63 | - | 8 x 10<sup>9</sup> | 20 |
| PA14 lysis 2 | 6.00 x 10<sup>8</sup> | 0.77 | - | 8 x 10<sup>9</sup> | 0.6 |
| PA14 lysis 3 | 5.40 x 10<sup>8</sup> | 0.67 | - | 8 x 10<sup>9</sup> | 0.2 |
| SA + phage | - | - | 0.34 | 8 x 10<sup>9</sup> | 0.2 |
| Poly lysis 1 | 6.40 x 10<sup>8</sup> | 0.81 | 1.14 | 8 x 10<sup>9</sup> | 2.5 |
| Poly lysis 2 | 3.50x 10<sup>8</sup> | 0.45 | 0.85 | 8 x 10<sup>9</sup> | 5 |

**Table 3:** training parameters for each model.

| Model | No. of training images | No. of PA14 cells | No. of MSSA476 cells | Epochs (training 1) | Epochs (training 2) |
| --- | --- | --- | --- | --- | --- |
| PA14 | 230 | 1600 | - | 200 | - |
| MSSA476 | 160 | - | 1300 | 200 | 50 |
| Bi-microbial | 480 | 2000 | 2200 | 150 | 80 |

The first model was trained for detecting the PA14 strain with rod shaped morphology (Figure 3-i). The dataset was generated by taking images from PA14 growth and lysis experiments. A total of 230 images containing over 1600 examples of PA14 morphology in different microscopic conditions (light intensity, gain, collimator position) were labelled manually by drawing bounding boxes. This dataset was then split into a ratio of 70:30 for training and validation respectively. The second model was trained detection of MSSA476 with cocci morphology or forming figures of 8 or aggregates with figures of 8 (Figure 3-ii). A total of 150 images containing over 1300 examples of MSSA476 morphology in different microscopic conditions were labelled and split into a 70:30 ratio like the PA14 model. The third model was trained for detection of both PA14 and MSSA476 cell strains. The dataset consisted of 480 images with over 2000 examples of PA14 morphology and 2200 examples of MSSA476 morphology. This dataset was generated by combining the first two datasets as well as adding examples from polymicrobial experiments where both PA14 and MSSA476 cells were present in one droplet.

**Figure 3:**
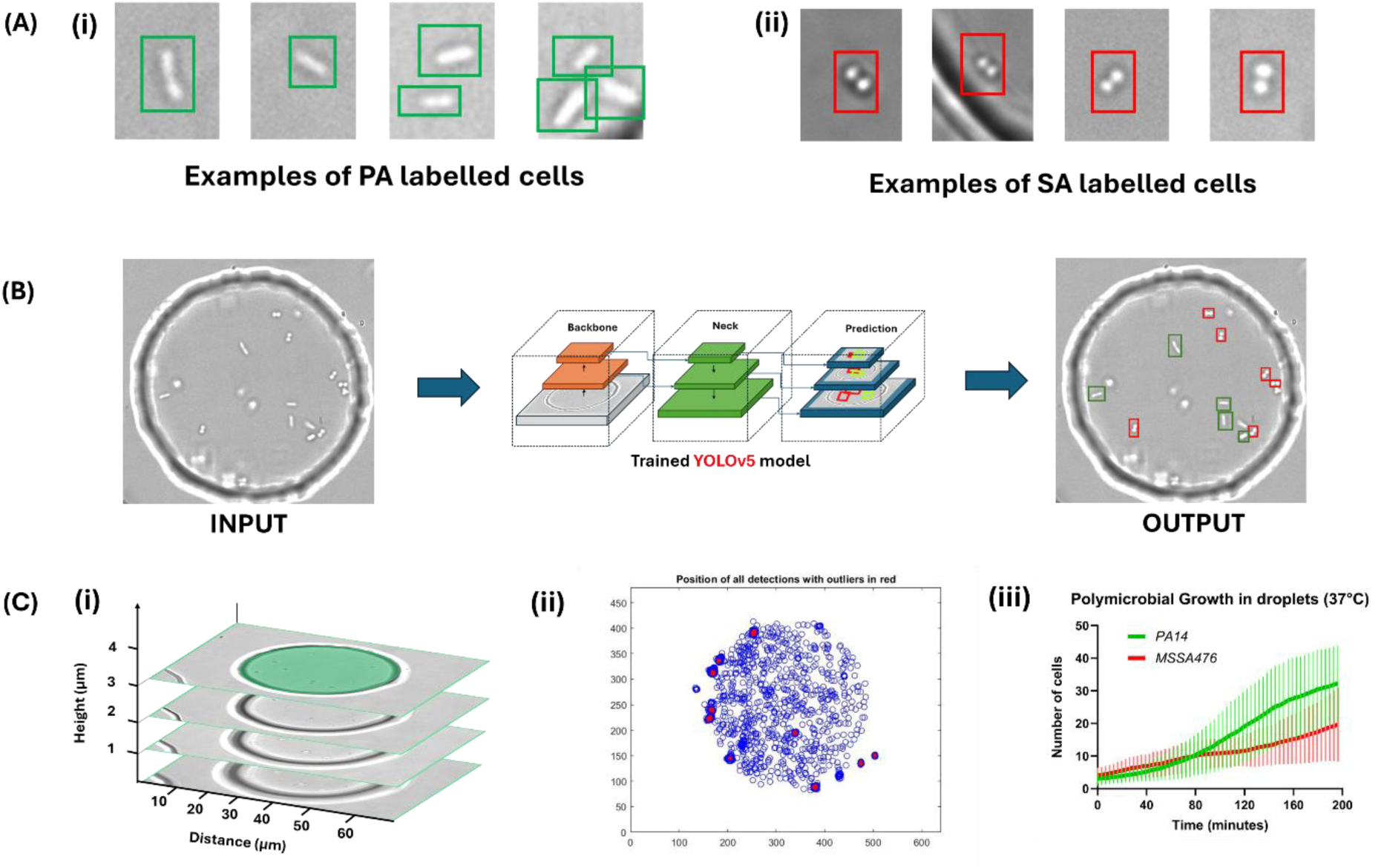
From labelling cells to generating accurate counts (A) The examples of PA14 and MSSA476 morphologies as labelled for training the YOLOv5 object detection model (A-i) Examples of cells with rod shaped morphology labelled as PA14 (A-ii) Examples of cocci-shaped cells labelled as MSSA476 (b) Object detection using YOLOv5x. Z-stack images are processed using the trained models to obtain bounding boxes with classification and positions of cells as seen in the output image. The green boxes are the model’s prediction for PA14 cells and the red boxes are predictions for MSSA476 cells (C) Generating cell counts (C-i) Each Z-stack is treated as one time point and cell count in obtained by superimposing all bounding boxes (C-ii) The blue dots represent the number of cells and their position detected during one experiment and the red dots represent the counts that were ignored due to clustering or false detections (x-axis is width in pixels and y-axis is height in pixels). (more information in the SI, section S3) (C-iii) Cell counts are obtained using moving average and plotted with standard deviation.

### 1.6 Two-species bacteria detection using YOLOv5

We developed a counting method to extract simultaneously the number of cells for PA and SA per droplet using Z-stack imaging. The Z-stack images collected using the imaging setup were utilized to perform morphology-based detections. Detection data from each image contained information on bounding box coordinates, detection scores and class information which were stored in corresponding detection files. All image processing was performed using MATLAB. For each time point, the corresponding images were loaded from a local folder, while the bounding box detections and class information were imported from detection files. Each Z-slice within a time point was analyzed for cell counts and classifications. Bounding box coordinates were calculated based on image dimensions (640×480 pixels). Cells were classified into PA or SA based on the class label and detection score. Only detections with confidence scores above a threshold of 40 % were considered valid to eliminate low-confidence detections.

To account for bacterial movement between slices, we calculated the Euclidean distance between bacteria across different slices. If two bacteria belonging to different Z-slices were detected within a predefined distance threshold (typically within a sphere of diameter 3 microns, which was determined using movement speed of the bacteria), one detection was excluded to avoid counting the same bacterium multiple times. In the case of 2 classes of cells, namely PA and SA, cells were detected accurately thanks to their marked difference in morphology. Occasionally, one cell was classified differently in two slices within the same stack, in which case the cell was classified based on the overall proportion of PA versus SA detections. Cells with a ratio over 50% of PA detections were labeled as PA, and vice versa. This classification method allowed real-time counting and localizing of PA and SA populations within droplets (Figure S3). Time assignment was performed by extracting the metadata from the first image in the sequence, which provided the initial time point. For subsequent images, we calculated the elapsed time in seconds relative to this initial time point using file metadata.

### 1.7 Cell doubling time calculation

Doubling times for bacterial populations were calculated by analyzing the slope of the bacterial growth curves. To ensure accuracy, we identified the portions of the plots with the steepest slopes within a 20-minute moving window. A MATLAB script was utilized to calculate the specific growth rates (SR) based on the selected regions. These specific growth rates (SR) were then used to derive the cell doubling times (*T*_*D*_) using the formula:

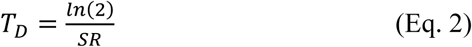

## 2. Results

We aimed at establishing a high-throughput method for the quantitative analysis of bacterial co-cultures and application to phage infection dynamics. The automated microscope stage enabled acquisition of Z stacks for up to 20 droplets over times of up to 24 hours, providing Z-stack time lapse imaging every 5 minutes. To correct for slight differences in focal planes for every droplet due to imperfect device flatness and possible vertical drift, we implemented a deep learning based autofocus method. Lateral drift was corrected by centering droplets using Hough transforms before every Z-stack imaging.

### 2.1 Deep learning for autofocus and bacterial cell detection

#### 2.1.1 Focus correction using image classification

We implemented an autofocus method using YOLOv8 based image classification. YOLOv8 is a version of the YOLO (You Only Look Once) real-time object detector which can perform tasks such as object detection and image classification. YOLOv8 was chosen due to its sophisticated data augmentation methods, which increases model robustness so it can be used on a wide range of imaging conditions, as well as its very high inference speed, which allows for very fast focal adjustments in-between imaging. A custom training dataset of 3320 images was assembled from previous microfluidics experiments using circular traps and Z-stack imaging. Images of differing quality were chosen with varying brightness, background noise and artifacts, as well as different trap and droplet sizes, bacterial species to maximize the robustness and flexibility of the model. These images were then split into 5 classes – traps or droplets that appeared ‘in focus’, slightly below or above the plane (approximately <3μm out of focus based on the Z-stack) and greatly out of focus above or below the plane (>3μm). The images were randomly split into a training and validation dataset, with an 85:15 ratio. The YOLOv8-nano model, chosen for its small size and therefore high detection speeds, was then trained on the custom dataset, which showed very high accuracy (∼96%) and low loss at the end of the training (refer to SI section 1 for more information on model training and accuracy parameters).

#### 2.1.2 Bacterial morphology detection using deep learning

*P. aeruginosa* strain *PA14 ΔflgK* and *S. aureus* strain *MSSA476* were chosen for this study based on their marked morphological differences (Figure 3A) and relevance to disease modelling (e.g., cystic fibrosis model). The flgK mutants had low motility compared to wild type PA14, and this simplified the data analysis given limited cell motion of the flgK mutants during Z-stack imaging.

All the data was generated in the form of brightfield Z-stack images corresponding to individual time points. A typical stack had 40 images imaged every 0.5 µm across a height of 20 µm. Every image was analyzed separately using single-stage object detection based on deep learning as mentioned in Section 1.5. YOLOv5x (extra-large version optimized for high accuracy) was the model of choice to perform the detection-based analysis. YOLOv5x architecture is made up of 25 convolutional layers, and better detection accuracy compared to other models in the YOLOv5 family ^27^. Three different models were trained for morphological detections for the analysis of three separate categories. The first model was trained to detect *P. aeruginosa* cells with rod-shaped morphology. The second model was trained to detect cocci shaped *S. aureus* cells with circular morphology. The third model was trained to detect both *P. aeruginosa* and *S. aureus* cells in a co-culture as two separate classes. The results of the model training are listed in Table 4. We achieved a detection accuracy of ∼99%, ∼91% and ∼97% for PA14 model, MSSA476 model and bi-microbial model respectively.

**Table 4:** mAP_0.5 values after model training (i.e. mAP at IOU 0.5 in section SI 1.2).

| Model | mAP_0.5 |
| --- | --- |
| PA14 | 99.4% |
| MSSA476 | 90.5% |
| Bi-microbial | 96.7% |

**Table 5.** Experimental conditions tested and deep learning models used for image analysis for mono and co-cultures of PA14/MSSA476 and exposure to P278 phage with fastest and average cell-doubling rates.

| Experiments | Species analyzed | Culture type | Fastest cell doubling rate (minutes) | Average cell doubling rate (minutes) | Detection model used |
| --- | --- | --- | --- | --- | --- |
| PA14 | PA14 | Monoculture | 26.52 | $33.96 \pm 6.69$ | PA14-v5 |
| MSSA476 | MSSA | Monoculture | 41.48 | $69.49 \pm 17.78$ | MSSA-v5 |
| MSSA476 + P278 | MSSA | Monoculture | 44.39 | $69.16 \pm 16.03$ | MSSA-v5 |
| PA14 with MSSA | PA14 | Co-culture | 30.96 | $35.055 \pm 4.86$ | Bimicrobial-v5 |
| MSSA with PA14 | MSSA | Co-culture | 43.39 | $73.34 \pm 21.07$ | Bimicrobial-v5 |
| MSSA+PA14+P278 (MOI 2.5) | MSSA | Co-culture | 60.21 | $100.69 \pm 30.08$ | Bimicrobial-v5 |
| MSSA+PA14+P278 (MOI 5) | MSSA | Co-culture | 80.74 | $123.28 \pm 37.113$ | Bimicrobial-v5 |

We generated anchored droplets using individual cell species PA14-flgK and MSSA476 in LB media as well as their co-cultures. Table 1 and 2 (methods) show all experimental conditions tested using our anchored droplet-based setup and the detection models used in all cases.

#### 2.1.3 Growth of individual bacterial species in anchored droplets

The first step in validating our droplet co-culture methodology was to conduct growth experiments for individual species within droplets. The time resolution was dependent on the total number of droplets screened and the number of images recorded per Z-stack. Typically, the platform would perform a screen of 10 droplets in about 1 minute. To correct for counting errors, we calculated a moving average of cell numbers obtained across a window of 8 consecutive time-points. Errors included missed detections close to the water-oil or trap boundary, difficulty in counting individual SA cells within aggregates. Static detections were removed as they were most likely false detections (see SI section 3). The PA14-v5 model built solely with images of PA14 cells was used to analyze experiments involving the PA14 cell strain including the droplet growth control as well as the interaction between PA14 cells and P278 phage. The MSSA-v5 model built with images of SA cells was used to analyze all MSSA476 related data including MSSA growth controls as well as the interaction between MSSA476 and P278 phage. The Bimicrobial-v5 model was used to analyze all co-culture experiments including co-culture growth and P278 interaction experiments.

#### 2.1.4 Growth of PA14 in anchored droplets at 37°C

Experiments were conducted to analyze the growth of PA14 cells in droplets starting at varying bacterial densities as mentioned in Table 1. Assuming an ellipsoidal droplet of ⁓11 pL in volume, the initial mean number of cells in each droplet could be anticipated with Poisson statistics based on the loading optical density (OD_600_) of the cell culture. Across different droplets, the data was analyzed using our cell counting method to calculate specific growth rates and cell doubling times as explained in methods. Figure 4A shows the change in cell count over time. For *P. aeruginosa* strain *PA14-flgK*, the fastest cell doubling time obtained using our method mentioned in Section 1.7, was 26.52 minutes with an average cell doubling time of 33.96 ± 6.69 minutes calculated over all droplets observed and analyzed.

**Figure 4:**
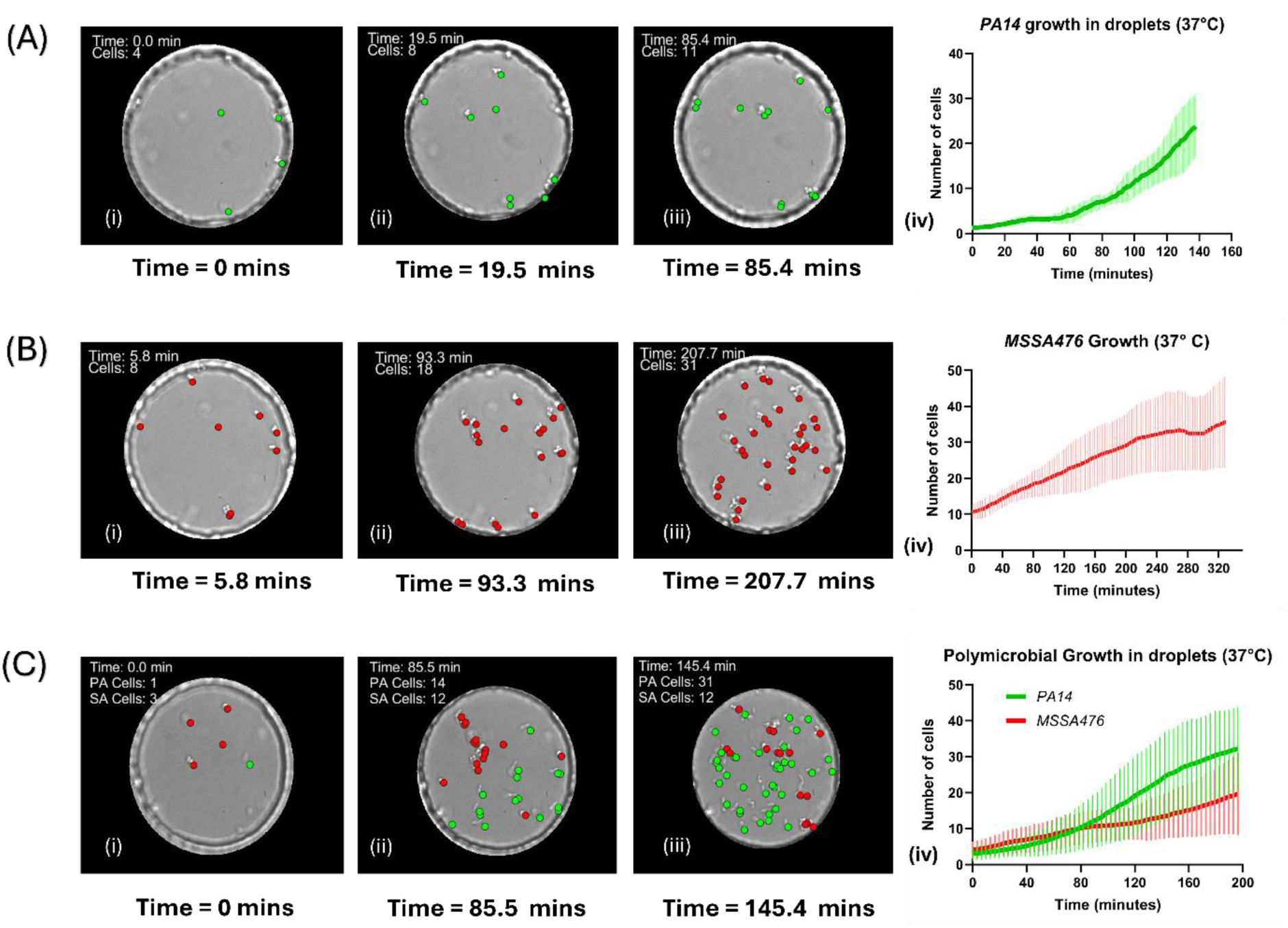
Growth dynamics of *Pseudomonas aeruginosa* PA14, *Staphylococcus aureus* MSSA476, and their polymicrobial interaction in droplets at 37°C. (A) Growth of *P. aeruginosa* PA14 over time in droplets at 37°C. The line plot shows the mean number of cells (solid green line) as a function of time, with the shaded area representing the standard deviation across growth experiments in droplets. Corresponding brightfield images depict snapshots of a selected droplet at various points, with individual detected cells marked by green dots. (B) Growth of *S. aureus* MSSA476 over time in droplets at 37°C. The red line and shaded region represent the mean and standard deviation respectively. Brightfield images highlight cell growth within a droplet at different time points, marked by red dots. (C) Polymicrobial growth of *P. aeruginosa* PA14 and *S. aureus* MSSA476 in droplets at 37°C. The green and red lines represent the mean cell counts of *P. aeruginosa* (PA) and *S. aureus* (SA), respectively, with shaded regions indicating standard deviations. Representative brightfield images illustrate the evolution of a co-culture within a droplet at different time points, with *P. aeruginosa* cells marked by green dots and *S. aureus* cells by red dots as seen in Movie S2.

#### 2.1.5 Growth of MSSA476 in anchored droplets at 37°C

For *S. aureus* strain MSSA476, the cell loading densities tested are mentioned in Table 1. The fastest cell doubling time observed was calculated as 41.48 minutes with an average cell doubling time of 69.49 ± 17.78 minutes. For one example droplet, Figure 4B shows the change in cell number over time.

#### 2.1.6 Co-culture growth of PA14 and MSSA476 in anchored droplets at 37°C

To generate droplets containing both PA14 and MSSA476 cells, cells of each species in exponential growth phase were mixed in specific ratio as detailed in Table 1. A total of 12 droplets were analyzed across two experiments. For PA14 cells in a co-culture with MSSA476 cells, the fastest cell doubling rate was calculated to be 30.96 minutes with an average cell doubling time of 35.06 ± 4.86 minutes. For MSSA476 cells growing in a co-culture with PA14 cells, the fastest cell doubling rate was calculated at 43.39 minutes with an average cell doubling rate of 73.34 ± 21.07 minutes. Starting at similar cell numbers, PA14 cells in most cases become the dominant species and outgrow MSSA476 cells. Individual droplet cell numbers can be seen in Figure S8 in SI.

### 2.2 Interaction of phage P278 with both species

To uncover bacteria-phage dynamics in a bi-species community, we tested a previously uncharacterized *P. aeruginosa* bacteriophage named P278 with both cell species individually and in co-cultures. Phage P278 was isolated by screening ⁓70 phages from the Exeter Phage Library against PA14-flgK. The result of the screen is shown in SI information section 2 Figure S5. Phage 278 was chosen for its good lytic efficiency against PA14-flgK and absence of apparent impact on SA growth.

Phage life cycle parameters were initially characterized using the one step growth curve method. The burst size (average number of phages released per infected cell) was calculated to be 46 and the time to first burst was calculated as 37 minutes (Figure S6). Experiments were performed at various initial multiplicity of infection (MOI) to understand the relationship between phage loading and individual and polymicrobial growth dynamics. MOIs were calculated for every experiment as described in Table 2.

#### 2.2.1 Interactions between PA14 and phage P278 in anchored droplets

Experiments were conducted at 3 different MOI conditions to observe the impact of lytic phage P278 on PA14-flgK cell strain. In all phage experiments, we targeted an initial cell count fewer than 15 cells from both species, such that they were still in exponential growth phase but were in sufficient numbers for quantifying decrease accurately.

The experimental conditions are listed in Table 2. Figure 5A represents the response of PA14 strain with an MOI of 0.6. A total number of 15 droplets were observed over 16 hours. Cells were found to initially grow for the first 50 minutes before lysing afterwards. The population declined significantly over the duration of the experiment, reaching close to zero PA cells after 8 hours. Figure 5B shows the lysis of PA14 cells at MOI of 20. A total of 8 droplets were observed for over 3 hours. We observed a change in PA14 cell morphology - sufficient for cells not to be detected by the model - within 20 minutes, indicating faster cell lysis due to excessively high number of phage particles compared to the number of cells in each droplet. To investigate low phage loads, we also conducted an experiment with an MOI of 0.2. This corresponded to 4-14 phages per droplet given to chosen bacterial densities. We imaged 12 droplets over 20 hours and observed an initial increase in the number of cells followed by a decrease and stabilization in cell number over time. Figure 5C represents the dynamics of the cell-phage interaction. Individual curves for all droplets are shown in SI figure S10.

**Figure 5:**
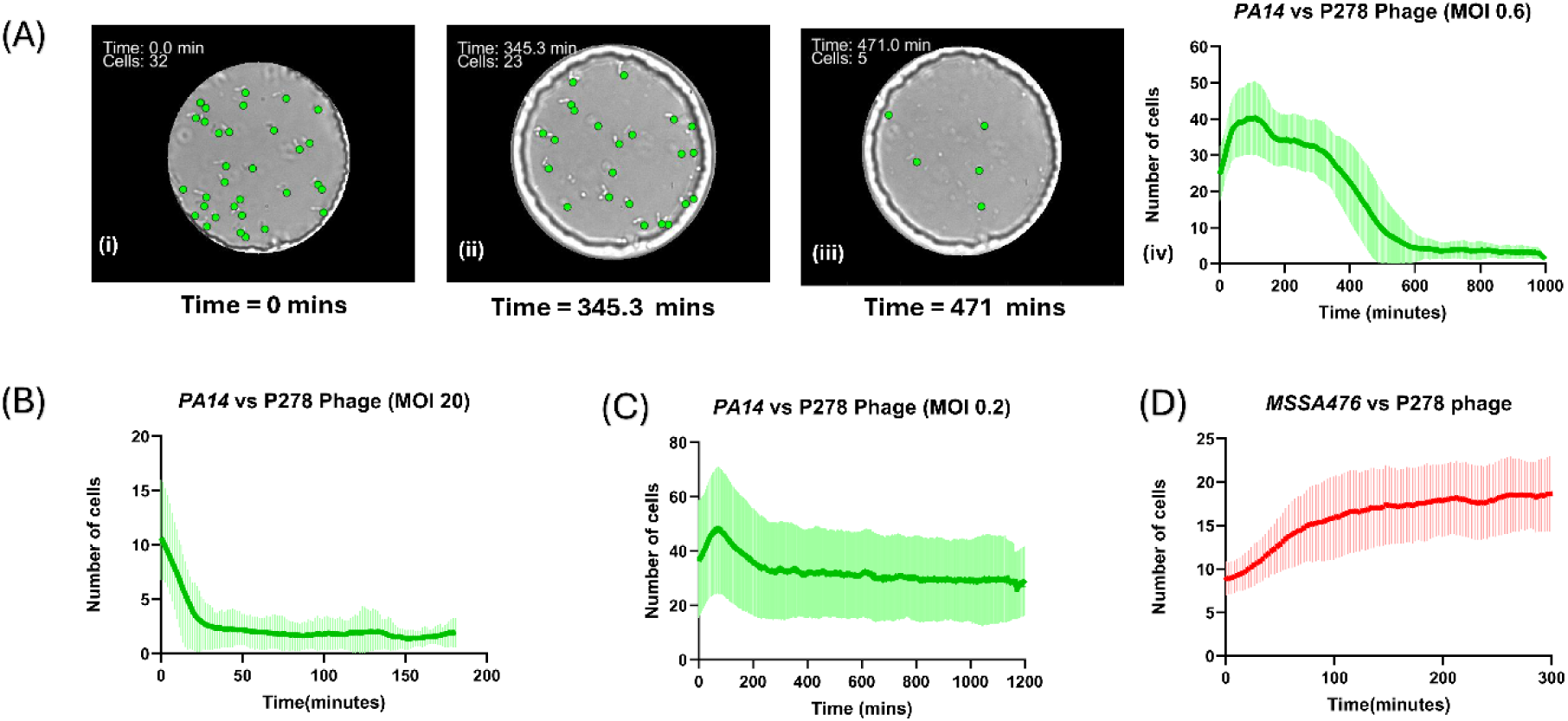
Effect of P278 phage on *Pseudomonas aeruginosa* PA14-flgK and *Staphylococcus aureus* MSSA476 populations at different multiplicities of infection (MOIs). Growth dynamics of PA14-flgK exposed to P278 phage at MOIs (A) Representative brightfield images for MOI 0.6 show droplets at various time points over 500 minutes, with *P. aeruginosa* cells marked by green dots cells. (B) Growth dynamics of PA14-flgK exposed to P278 phage at MOI 20 (c) Growth dynamics of PA14-flgK exposed to P278 phage at MOI 0.2. The line plots show the mean cell count over time (solid green line), with the shaded area indicating the standard deviation. (D) Growth of *S. aureus* MSSA476 in the presence of P278 phage. The red line represents the mean cell counts over time, with the shaded area showing standard deviation. Movie S3 shows the lysis of PA14 cells in droplets.

#### 2.2.2 Interactions between MSSA476 and phage P278 in anchored droplets

We conducted control experiments to check if phage P278 displayed any interaction with MSSA476 cells in droplets. Figure 5D represents the growth observed when phage P278 was introduced to MSSA476 cells in an exponential growth phase. The red line represents the average growth, and the shaded region represents the standard deviation. A total of 6 droplets were observed across 3 experiments. The fastest cell doubling rate was calculated to be 44.39 minutes with an average cell doubling rate of 69.16 ± 16.03 minutes.

#### 2.2.3 Interactions between co-cultures of PA14 and MSSA476 with phage P278 in anchored droplets

The final set of experiments were conducted to study the impact of phage P278 on a co-culture of PA14 and MSSA476 cells. Two experiments were conducted at MOI of 2.5 and MOI of 5 respectively. Figure 6A represents the interaction between PA14 and MSSA476 strains in the presence of pseudomonas phage P278. The red and green lines represent the average response of MSSA476 and PA14 strains respectively, with the red and green shaded region showing the corresponding standard deviations. In the first experiment with MOI 2.5, over 10 droplets were observed for over 8 hours. The presence of phage P278 was found to suppress the growth of PA14 cells long-term with cell numbers staying at low levels (<5) over the 8 hours duration of the experiment. The fastest cell doubling rate for MSSA476 was calculated as 60.21 minutes with an average cell doubling rate of 100.69 ± 30.08 minutes. At MOI 5,8 droplets were monitored and the fastest cell doubling rate for MSSA476 was calculated at 80.74 minutes with an average cell doubling rate of 123.28 ± 37.113 minutes.

**Figure 6:**
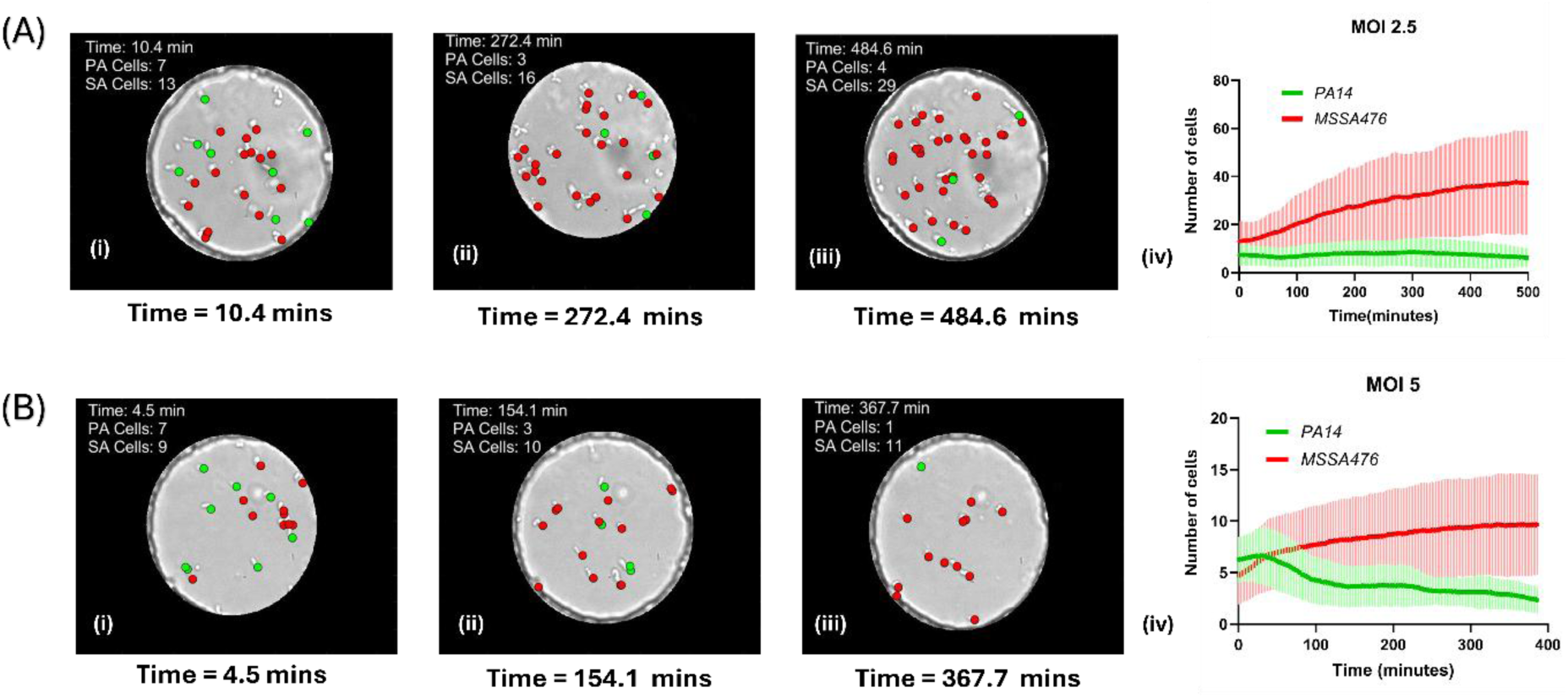
Effect of P278 phage on polymicrobial populations of *Pseudomonas aeruginosa* PA14 and *Staphylococcus aureus* MSSA476 at MOI 2.5 as seen in Movie S4. (A) Polymicrobial growth dynamics of *P. aeruginosa* PA14 (green) and *S. aureus* MSSA476 (red) in the presence of P278 phage at MOI 2.5. (Ai-Aiii) Brightfield images at different timepoints highlight the reduction in *P. aeruginosa* cells over time, with S. aureus continuing to grow. (A-iv) Line plot shows the mean cell count over time for both species, with shaded areas representing standard deviations. (B) Polymicrobial growth dynamics at a higher P278 phage MOI of 5. (Bi-Biii) Brightfield images at different timepoints highlight the reduction in *P. aeruginosa* cells over time, with S. aureus continuing to grow. (B-iv) Line plot shows the mean cell count over time for both species, with shaded areas representing standard deviations.

## 3. Discussion

We have established a method for the quantitative analysis of bacteria-phage interactions within model bi-microbial communities in a label-free format. This work relies on the confinement of bacterial communities within stable microdroplets and the ability of deep learning models to learn specific cell morphologies. Further, automation of the imaging platform enabled acquisition of large datasets for both technical and biological repeats. Specifically, this was achieved thanks to the implementation of a novel deep learning-driven object detection technique to achieve reproducible imaging over long durations tested up to 24 hours.

We demonstrate precise quantification of growth and interactions between bacteria and bacteriophages within monocultures and co-culture micro-environments. Our method of combining anchored microfluidic droplets with object detection-based image analysis provides several advantages compared to traditional microbiology assays. Unlike absorbance-based liquid (batch) co-culture experiments, which cannot distinguish between different bacterial species, our label-free imaging approach enables accurate differentiation and counting by exploiting morphological differences between species. This eliminates the need for repeatedly plating cells on selective agar plates, or performing other secondary assays, greatly reducing time and resources needed to perform bi-microbial studies.

It is important to maintain focus throughout the duration of experiments to ensure the Z-stack images are taken across the whole depth of the droplets. In this study, we have utilized deep learning-based image classification with 5 classes corresponding to 5 different focal ranges to counter the effects of systematic device tilt and unpredictable focal changes due to constant movement of the automated stage as well as other external factors including table vibrations. The speed of focal adjustment is important to minimise delays, acquire more time points per droplet and therefore obtain more accurate cell counts. Therefore, YOLOv8n (nano) was chosen for image classification. The integration of YOLOv8 for focus correction significantly improved image acquisition efficiency and quality. The model achieved a 96% classification accuracy using the training dataset facilitating robust imaging conditions as well as precise focal adjustments. We conducted experiments screening or droplets for over ∼20 hours with successful integration of this autofocus functionality (Supplementary Figure S1).

For bacterial detection, we focused on morphology-based analysis as maintenance of cell morphology (especially shape and size) is closely linked to their fitness ^32^. In particular, cells lysed by phages undergo dramatic morphological changes. This limits applicability to species with marked differences in morphology caused by interactions with antimicrobials such as phages.

In this study, we have employed YOLOv5x (extra-large) as it offers several advantages over previous versions of YOLO: less training time (∼4x faster training), better mAP precision, ease of implementation, reduced model complexity and transfer learning. Three different models trained on PA14, MSSA476, and bi-microbial datasets achieved detection accuracies of ∼99%, ∼91%, and ∼97%, respectively, facilitating accurate identification bacterial morphologies across different experimental conditions. There remain errors in false positive detections such as dust and traps imperfections and false negative detections for which cells are missed. This occurred frequently when cells were located close to the trap and droplet boundary. In bi-microbial communities, every cell was detected multiple times, and we used the average class to minimise misclassification. In our study, errors in absolute cell counts were reduced through computing a moving average. We also excluded detections which did not move across time series, as they most likely corresponded to false detections.

For PA14 monocultures, the average doubling time was 33.96 ± 6.69 minutes. This matches literature values for cell doubling times observed in liquid culture assays for the parental strain ^33^. MSSA476 alone exhibited slower growth compared to PA14, with an average doubling time of 69.49 ± 17.78 minutes. This time was longer than reported values of e.g. 20 minutes when *S. aureus* was cultured in rich medium ^34^. These findings highlight intrinsic growth rate differences in the droplet format, likely influenced by metabolic disparities and environmental adaptation. In co-culture droplets, however, PA14 consistently outcompeted MSSA476, demonstrating a faster average doubling rate (35.055 ± 4.86 minutes) compared to MSSA476 (73.34 ± 21.07 minutes) and final mean yield ratio of 2.4 ±1.6 PA:SA. This suggests a competitive advantage for PA14, that has been well reported in literature ^10^. However, MSSA476 sometimes outcompeted PA14 (i.e. in approximately 10% of our experiments, Supplementary Figure S9), indicating that the droplet format is more conducive to balanced growth for the two species compared to traditional formats. This may be due to higher oxygen availability as the oil phase dissolves over 10 times more oxygen than water ^12,28^. Similar cell doubling rates were observed for PA14 cells in monoculture as well as in a co-culture with MSSA476 cells. However, cell doubling rates for MSSA476 in a co-culture were slightly higher compared to when MSSA cells grew in a monoculture.

In droplet experiments, we observed that lysis by P278 phage caused an initial morphology to change for PA14 cells followed by progressively degradation. Phage P278 exhibited strain-specific lytic activity within droplets. PA14’s growth was significantly suppressed across all MOI conditions, with rapid lysis observed at MOI of 20 indicating that higher MOI leads to faster lysis of cells. The initial increase in cell number in experiments with MOI 0.6 and 0.2 can be attributed to the latent period being longer than cell division rate for PA14 in a monoculture. The fastest cell doubling rate for PA14 cells in a monoculture (∼27 minutes) was found to be lower than the time to lysis as observed in the one step growth curve (∼37 minutes). In contrast, MSSA476 was not affected by P278, maintaining growth rates comparable to phage-free conditions (∼70 minutes in both cases). In co-culture experiments, phage P278 selectively inhibited PA14, allowing MSSA476 to outcompete PA14. At MOI 2.5 and MOI 5, MSSA476 displayed long doubling times of 100.69 ± 30.08 minutes and 123.28 ± 37.113 minutes, respectively. We hypothesize that these extended cell doubling rates could be due to the presence of PA14 cell lysate and high numbers of phage particles within each droplet.

The droplet format helps uncover the heterogeneity of growth and responses to phage exposure. Thanks to stochastic cell loading, by targeting an average number of 10 cells per droplet, some droplets contained as little as one bacterium. The diversity of growth curves is illustrated in the individual plots shown in SI and can help shine light on assay reproducibility and the statistical significance of results. This enables the study the inoculum effects, i.e. the influence of initial bacteria densities on the efficacy of phage treatment. For example, the final yield for PA14 with P278 at MOI 0.2 was positively corelated with the initial cell number. Likewise, phage MOIs represent an average number for phage dosing, and absolute numbers of phages in each droplet will vary. In all phage experiments, we did not observe droplets in which cells did not undergo lysis, indicating all droplets had sufficient number of phages to initiate infection cycles. Unsurprisingly, we observed a dose-dependence between different MOIs but also noticed differences within the same MOIs tested. In general, the higher the MOI, the faster PA cell populations declined.

## 4. Conclusions

The deep learning bacterial detection framework enabled us to differentiate PA14 and MSSA476 in a mixed confined 3D environment based on morphological features. This capability is critical for understanding species-specific dynamics and interactions. The ability to observe and quantify bacterial dynamics in real time provides invaluable insights into growth patterns, competition, and antimicrobial efficacy.

The microfluidics platform’s adaptability to various experimental conditions (temperature, cell type and numbers), combined with automated imaging and analysis, enhances experimental throughput (currently up to 20 droplets imaged every 2 minutes) and reproducibility. The presented methodology can be extended to study other microbial and phage systems, enabling a deeper understanding of pathogen dynamics and co-evolution mechanisms. Additionally, it provides a robust platform for testing new antimicrobials and phage therapies in in-vitro polymicrobial settings.

Our approach will be also useful to study the dynamics of bacterial communities in response to external physical, chemical and biological stressors. In future, other phenotypes could be studied in parallel to cell localization such as morphological adaptation in response to stress or change in motility patterns^35,36^.

By overcoming the drawbacks of traditional microbiological methods, requiring repeated plating experiments or lengthy molecular assays over long timescales, our strategy offers a distinct framework for examining intricate microbial ecosystems with rapid evaluation of bacteria-phage dynamics. Future work could include the study of larger multi-species communities. This could be done by integrating additional imaging modalities (e.g., using fluorescent strains) and expanding deep learning models to include other bacterial species and morphological variants. This will further enhance the versatility of the analytical framework. Insights into the competitive dynamics of PA14 and MSSA476, particularly in the presence of phages, have implications for understanding microbial interactions in clinical settings such as cystic fibrosis infections. The findings highlight the potential of phages as targeted antimicrobial agents, particularly in polymicrobial infections where long-term *Pseudomonas* growth suppression is critical to outcomes in patients with chronic infections.

## Acknowledgements

The authors thank Dr. Nela Nikolic, Dr. Remy Chait, Dr. Tobias Bergmiller, Dr. Vasileios Anagnostidis for useful discussions and comments, Prof. Ben Temperton for access to the Exeter Citizen Phage Library, Prof. Stefano Pagliara for the bacterial strains. We acknowledge the LSI Technical Services Team at the University of Exeter and use of the Exeter Microfluidics Facility and Savchenko Centre for Nanoscience. For the purpose of open access, the author has applied a ‘Creative Commons Attribution (CC BY) licence to any Author Accepted Manuscript version arising from this submission.

## Data availability statement

Code used for YOLOv5 and YOLOv8 training are available on https://github.com/ultralytics.

All the MATLAB scripts used for analysis in this paper and Python code for Z-stack acquisition and drift correction are available at:

https://github.com/GielenLab/YOLOv5-v8-polymicrobial

Training images and model weights data will be shared upon reasonable request.

## 1. YOLO training for both autofocus and morphology detections

### 1.1 Deep learning for autofocus

An initial training of the image classification model using YOLOv8-Nano for autofocus u was used with a set of 300 images trained for 160 epochs. 5 successive training iterations of 55 epochs each were then carried out, with additional training images from model failures added each time, making the model more robust in its final iteration.

**Figure S1:**
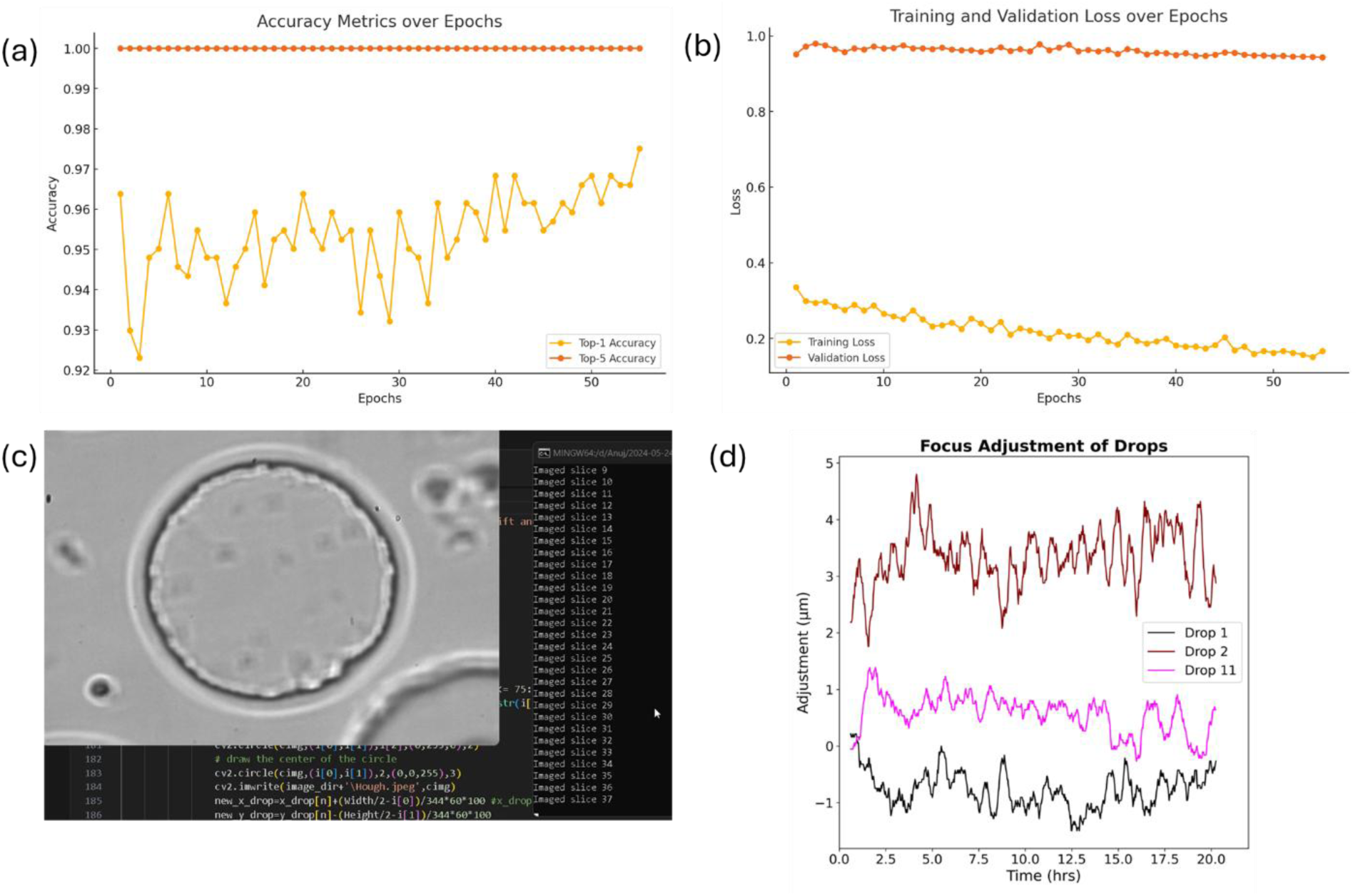
(a) Accuracy metrics over the last 55 epochs of training for image classification models for autofocus (b) Training and validation loss over the last 55 epochs for image classification training for autofocus model (c) Example of focus autocorrection during the experiment (d) Example of Z axis adjustments per droplet to correct focal drift during data acquisition experiments over 20 hours.

Top 1 classification accuracy is the measure of the percentage of times the model’s single highest-probability prediction (the “top 1” prediction) matches the true label. It is calculated as:

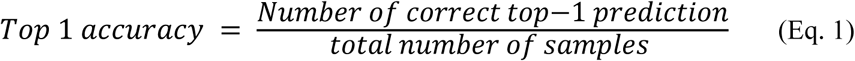

In our case, the image classification model using YOLOv8-Nano reached a top 1 accuracy of **97.51%** over the training as seen in Figure S1(a). Since there were only 5 classification classes, the top 5 accuracy remained 100% over the 50 epochs as seen in Figure S1(a). The final training loss was observed to be 0.16 and final validation loss reached a value of 0.94 indicating strong model performance with robust learning and accurate predictions.

### 1.2 Deep learning for cell morphology detection

For YOLOv5, precision and recall are calculated using standard definitions from the field of object detection. Precision is the fraction of true positive detections out of all positive detections made by the model. Mathematically, it can be expressed as:

**Precision = True Positive / (True Positive + False Positive)**
where True Positive is the number of correctly predicted objects, and False Positive is the number of objects predicted by the model that do not exist in the ground truth data.

Recall is the fraction of true positive detections out of all ground truth positives. Mathematically, it can be expressed as:

**Recall = True Positive / (True Positive + False Negative)**
where True Positive is the number of correctly predicted objects, and False Negative is the number of objects in the ground truth data that were not detected by the model. Other important measures for measuring the performance of a trained model are training loss and validation loss. IOU is defined as the ratio of the intersection area between the predicted bounding box and the ground truth bounding box to the union area of these two boxes. It is expressed mathematically as:

**IOU = (Area of intersection) / (Area of union)**

Mean average precision(mAP) values are measured as a function of IOU values. mAP_50 refers to the precision of detection at 50% IOU threshold.

While the general trend for a well-trained object detection model for precision and recall is an increase close to 1 over the training epochs, the trend for training and validation loss is a decrease towards 0. For each model, the training conditions are mentioned in each section below.

#### 1.2.1 PA14 morphology detection model

The dataset was generated by taking images from PA14 growth and lysis experiments. A total of 230 images containing over 1600 examples of PA14 morphology in different microscopic conditions (light intensity, gain, collimator position) were labelled manually by drawing bounding boxes. This dataset was then split into a ratio of 70:30 for training and validation respectively. For PA14 model, even without the transfer learning, we reached a mAP_50 value of 99.5%. The total precision and recall values at the end of the training were 0.99 and 0.98 respectively. The final values of training and validation loss were 0.0146 and 0.008 demonstrating effective training. The training metrics are shown in Figure S2.

**Figure S2:**
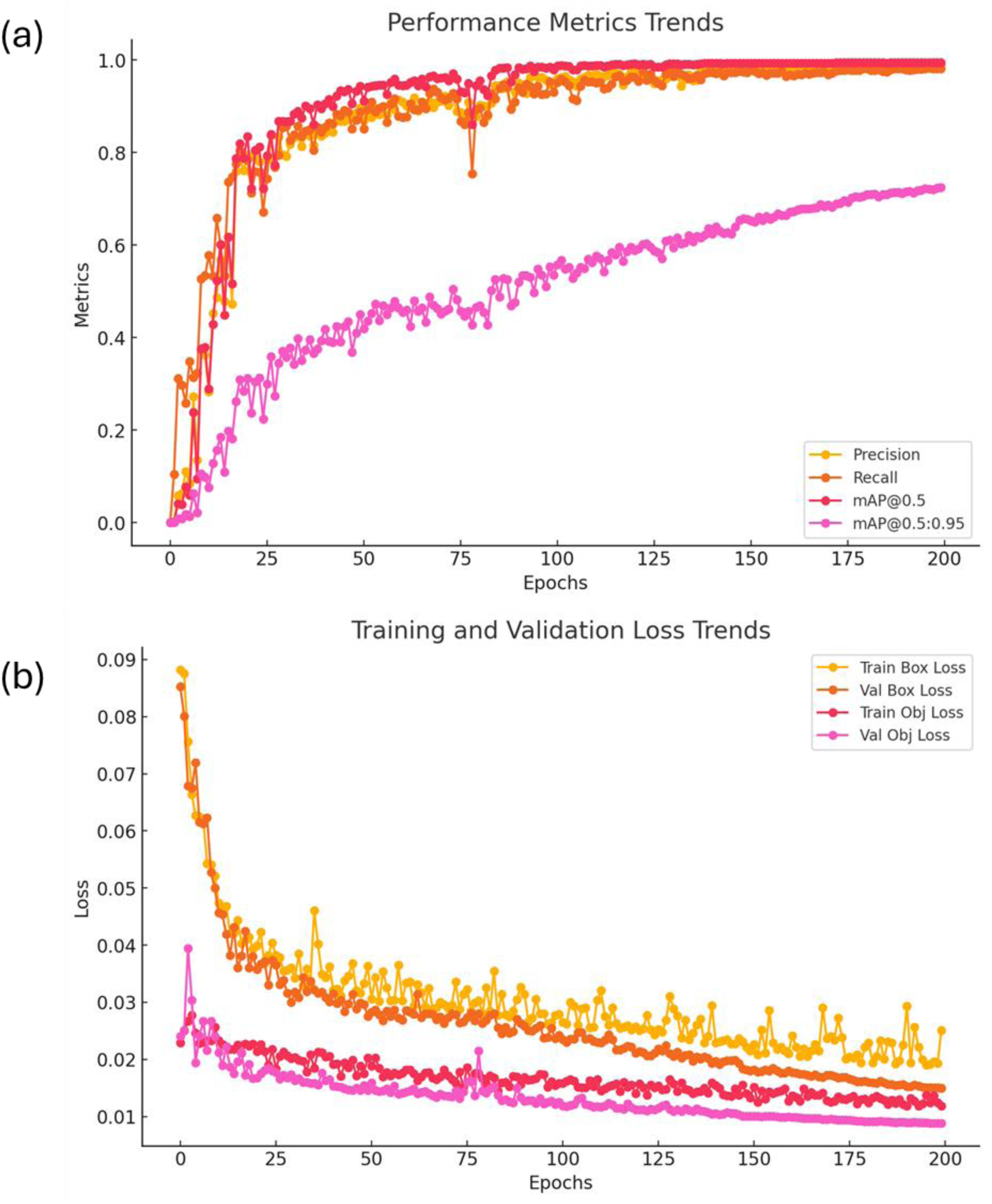
Training metrics for PA14 morphology detection model.

#### 1.2.2 MSSA476 morphology detection model

A total of 150 images containing over 1300 examples of MSSA476 morphology in different microscopic conditions were labelled and split into a 70:30 ratio like the PA14 model. The training was performed in two iterations using transfer learning. The original model was trained for over 200 epochs after freezing the first 10 layers of the model. The mAP_50 obtained at the end of this step was 83.6% with training and validation loss at 0.025 and 0.019 respectively as seen in Figure S3(a) and S3(b). Once the training finished, the best weights were obtained from the first training and retrained for 60 epochs after unfreezing the first 10 layers to improve mAP values. After the final training with the transfer learning step, the final mAP_50 was 90.5% with the training and validation loss values at 0.023 and 0.017 as seen in Figure S3(c) and S3(d). This model was used for detection of MSSA476 cocci-shaped morphology.

**Figure S3:**
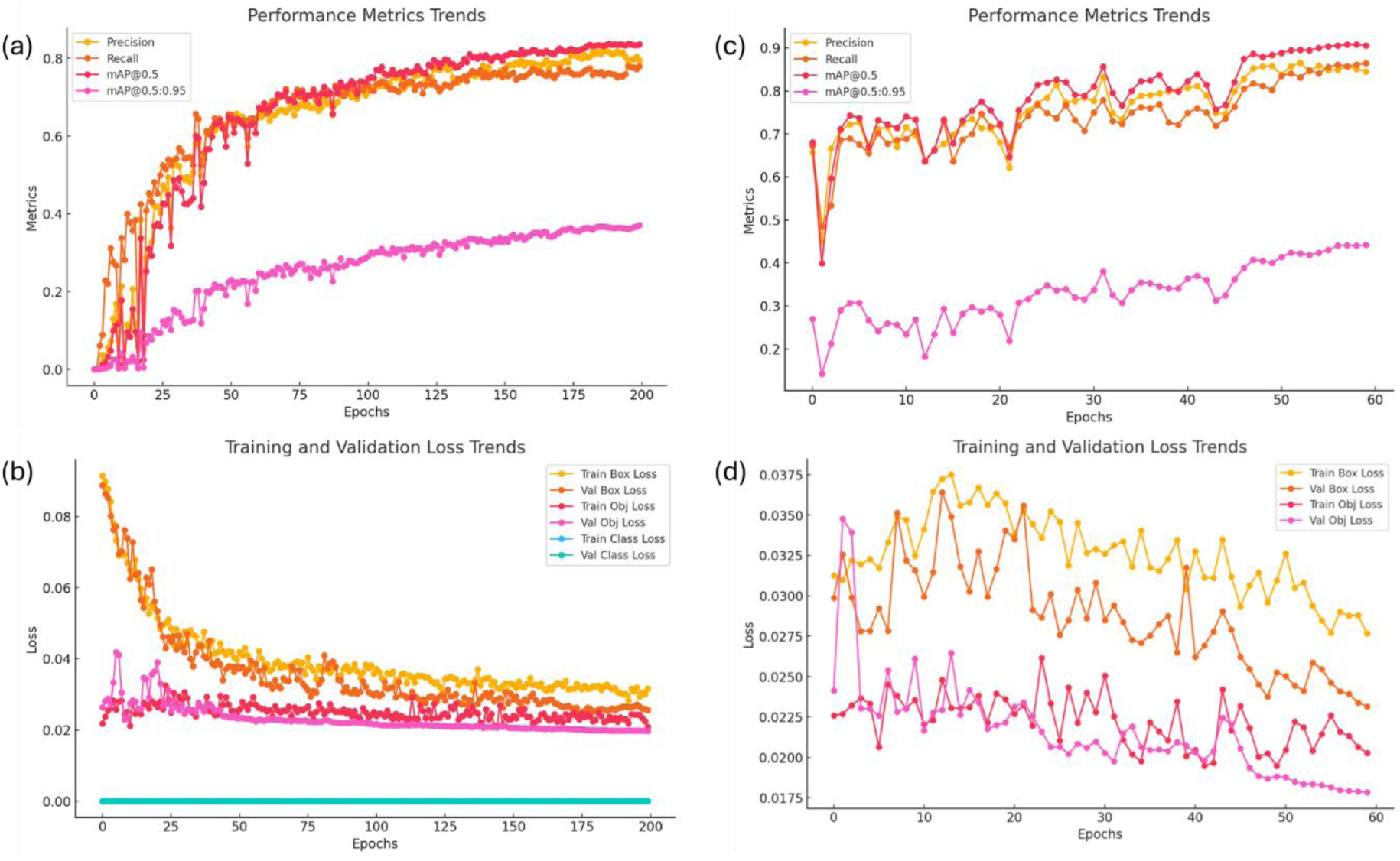
Training metrics for MSSA476 morphology detection. (a-b) Training metrics for the first 200 epochs where the first 10 layers of the model were frozen (c-d) Training metrics after obtaining weights from the first training iteration and retraining them after unfreezing the layers of the model.

#### 1.2.3 Polymicrobial model to detect both PA14 and MSSA476 using a single model

The third model was trained for detection of both PA14 and MSSA476 cell strains. The dataset consisted of 480 images with over 2000 examples of PA14 morphology and 2200 examples of MSSA47 morphology. This dataset was generated by combining the first two datasets as well as adding examples from polymicrobial experiments where both PA14 and MSSA476 cells were present in one droplet. Similar to the MSSA476 model, this model was also trained using transfer learning. The first iteration of training was done for 150 epochs after freezing the first 10 layers of the model. The mAP value obtained was 91.5% with a training and validation loss of 0.016 and 0.001 as seen in Figure S4(a) and S4(b). The transfer learning involved training for another 60 epochs after unfreezing the first 10 layers. This improved the mAP value to 97% with a training and validation loss of 0.013 and 0.0005 as seen in Figure S4 (c) and S4 (d).

**Figure S4:**
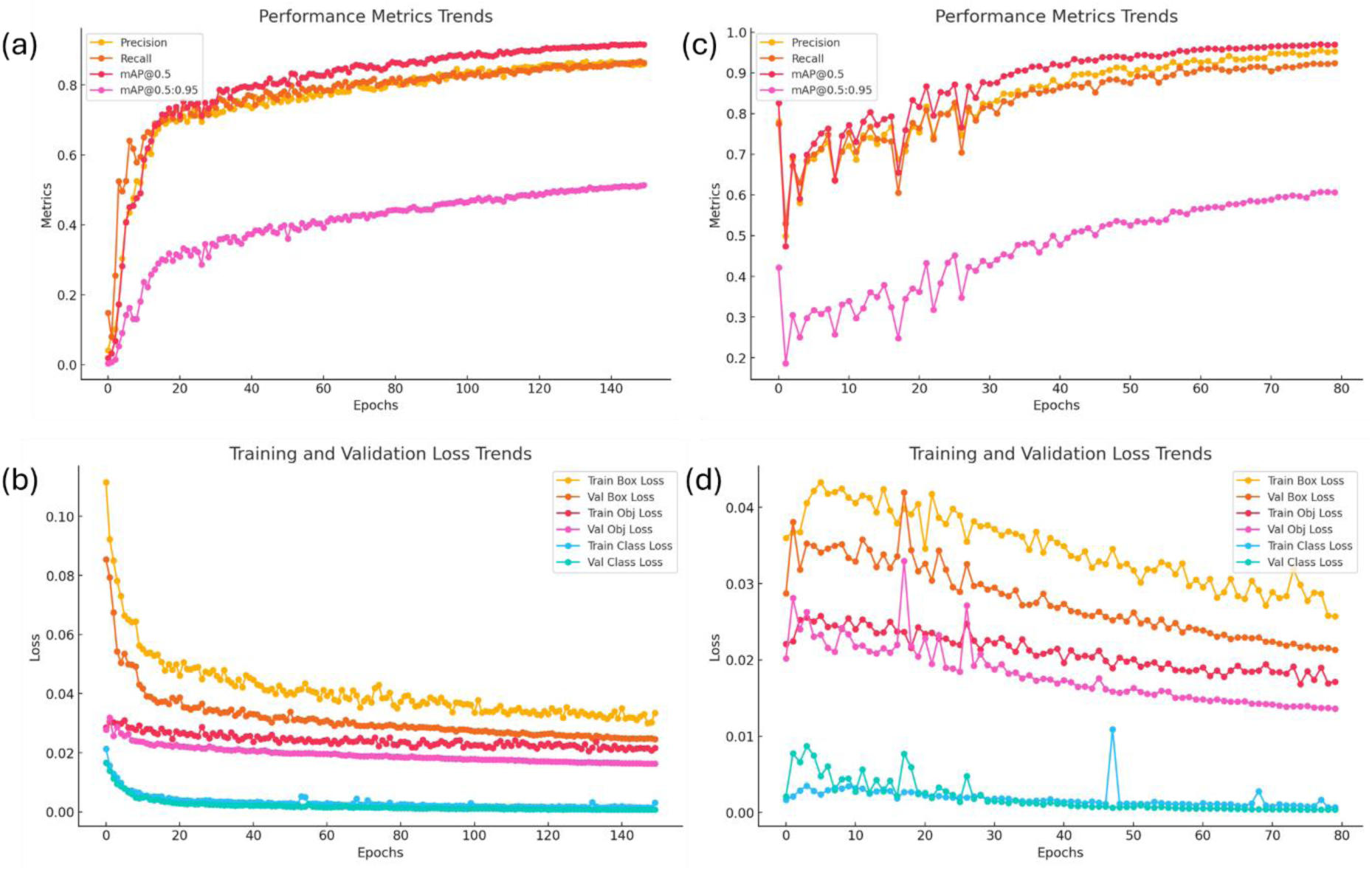
Polymicrobial model training with transfer learning. (a-b) Training metrics for the first 150 epochs where the first 10 layers of the model were frozen (c-d) Training metrics after obtaining weights from the first training iteration and retraining them after unfreezing the layers of the model.

## 2. One-step growth curve for phage P278

**Figure S5:**
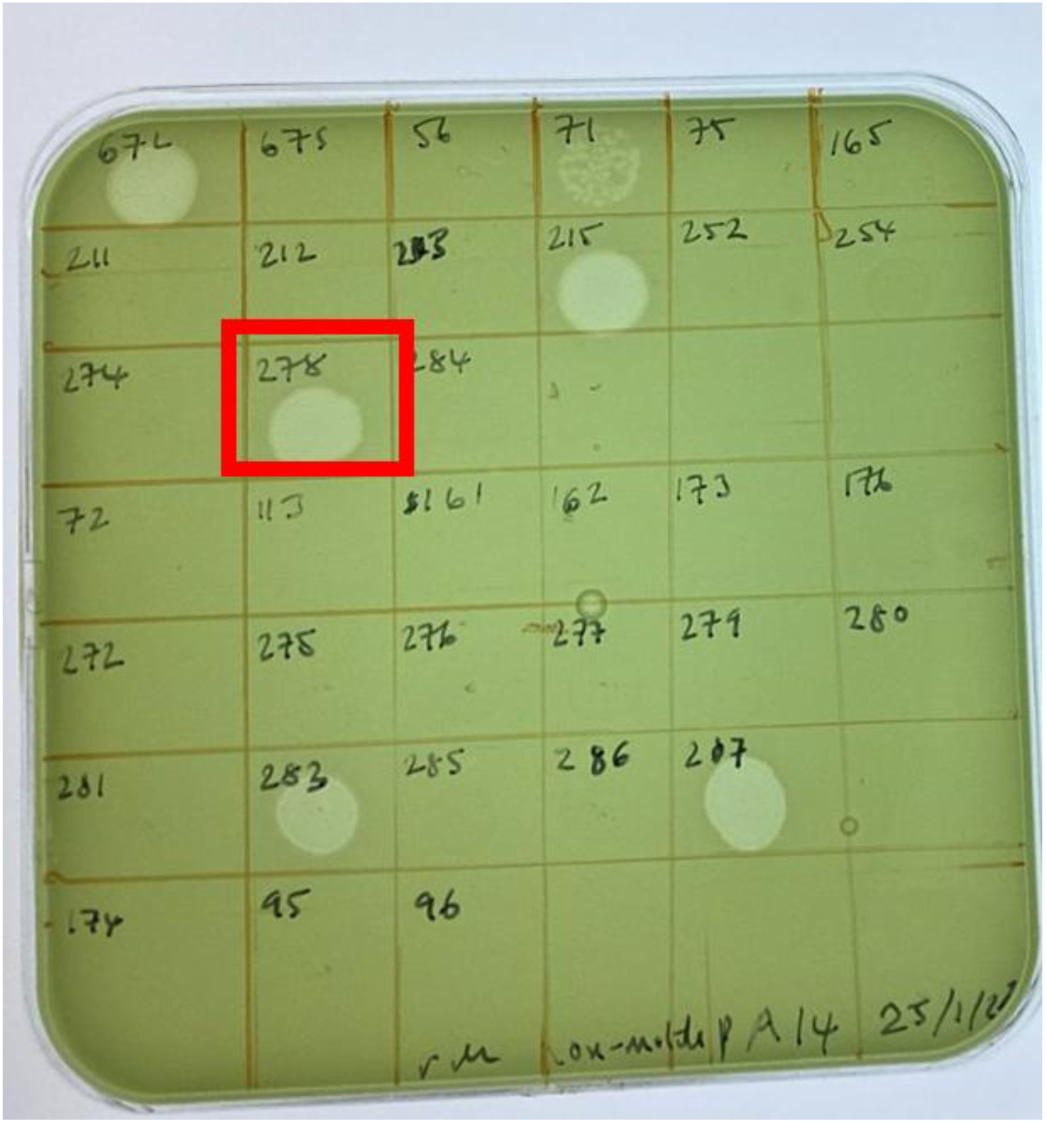
Characterizing different phages against PA14-flgK strain P278 was chosen as it formed clear plaques on the top agar bed with PA14-flgK cells.

To characterize phage P278 for time to lysis and burst size, we performed a standard one-step growth curve assay. A single colony of PA14-flgK cells was grown in 5 mL of LB media overnight. 50 µL of cells from overnight culture were diluted into 5 mL of fresh LB media. The cells were grown up to an OD_600_ of 0.650 corresponding to 6 x 10^8^ CFU/mL. The phage titer for P278 had a concentration of 8 x 10^9^ PFU/mL. 24 Eppendorf tubes with volumes 1.5 mL were autoclaved and prepared for the experiment. Each Eppendorf tube would contain 5 µL of phage titer, 332 µL of cells at OD 0.65 and 663 µL of fresh LB bringing the volume to a total of 1 mL. This would result in an MOI of 0.2 in each tube. All 24 tubes were placed on a heating block set at 37 °C, shaking at 200 RPM. We chose to sample 8 points over the duration of 1 hour; each point measured every 8 minutes with 3 repeats. For each point, 200 µL of solution was pipetted into a 96 well filter plate and centrifuged at 1200 g for 2 minutes to separate out the free phage left in the solution. After repeating this process across 8 time-points over 56 minutes, a 10-fold dilution series in SM media was performed for each point 8 times.

LB agar-based Petri dishes were prepared to perform the plaque forming assay to calculate the number of free phages for each time point. 100 µL of PA14 cells at OD 0.7 were mixed with 5 µL of CaCl_2_ and MgCl_2_ and 5 mL of soft agar and poured on top of each petri dishes with LB agar base to create a bed of *PA14* cells as a top layer. Using a multichannel pipette, 2.5 µL of each concentration from the dilution series for each point was plated onto the Petri dishes and left overnight to calculate the concentration of free phage as seen in Figure S5. The resulting number of plaque-forming units (pfu) are listed in Table 1.

**Figure S6:**
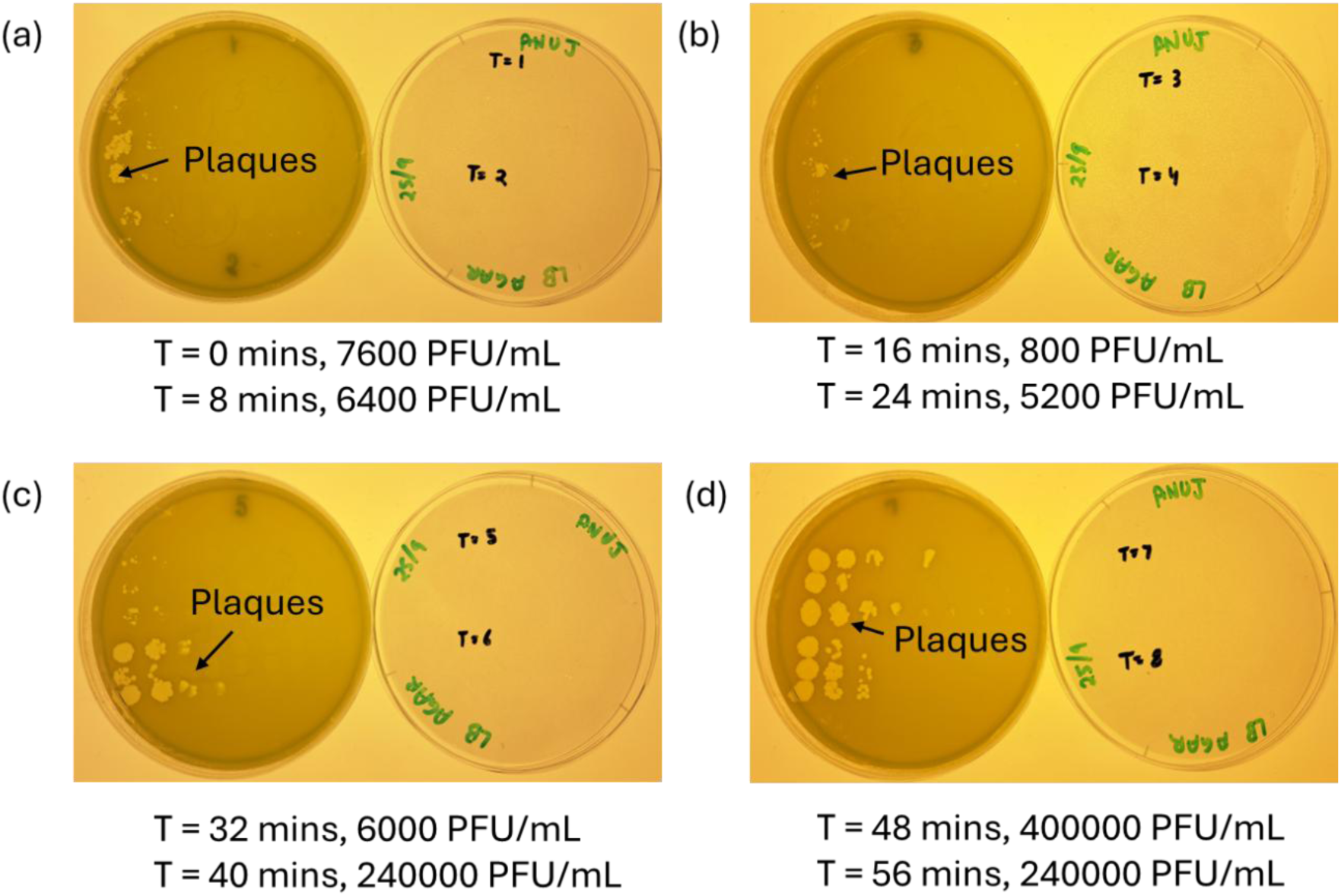
One-step growth curve assay to characterize P278 phage and its interaction with PA14 cells. (a) Time points 1 and 2 with dilution series and plaques formed. (b) Time points 3 and 4 with dilution series and plaques formed (c) Time points 5 and 6 with dilution series and plaques formed. (d) Time points 7 and 8 with dilution series and plaques formed.

**Table 1:**
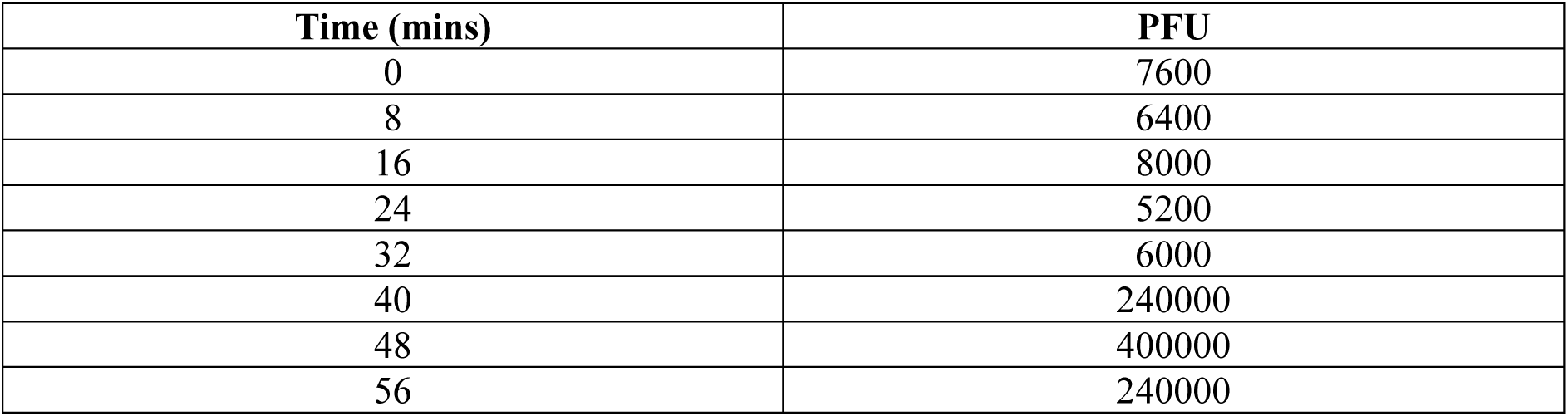
PFU count overtime points.

| Time (mins) | PFU |
| --- | --- |
| 0 | 7600 |
| 8 | 6400 |
| 16 | 8000 |
| 24 | 5200 |
| 32 | 6000 |
| 40 | 240000 |
| 48 | 400000 |
| 56 | 240000 |

**Figure S7:**
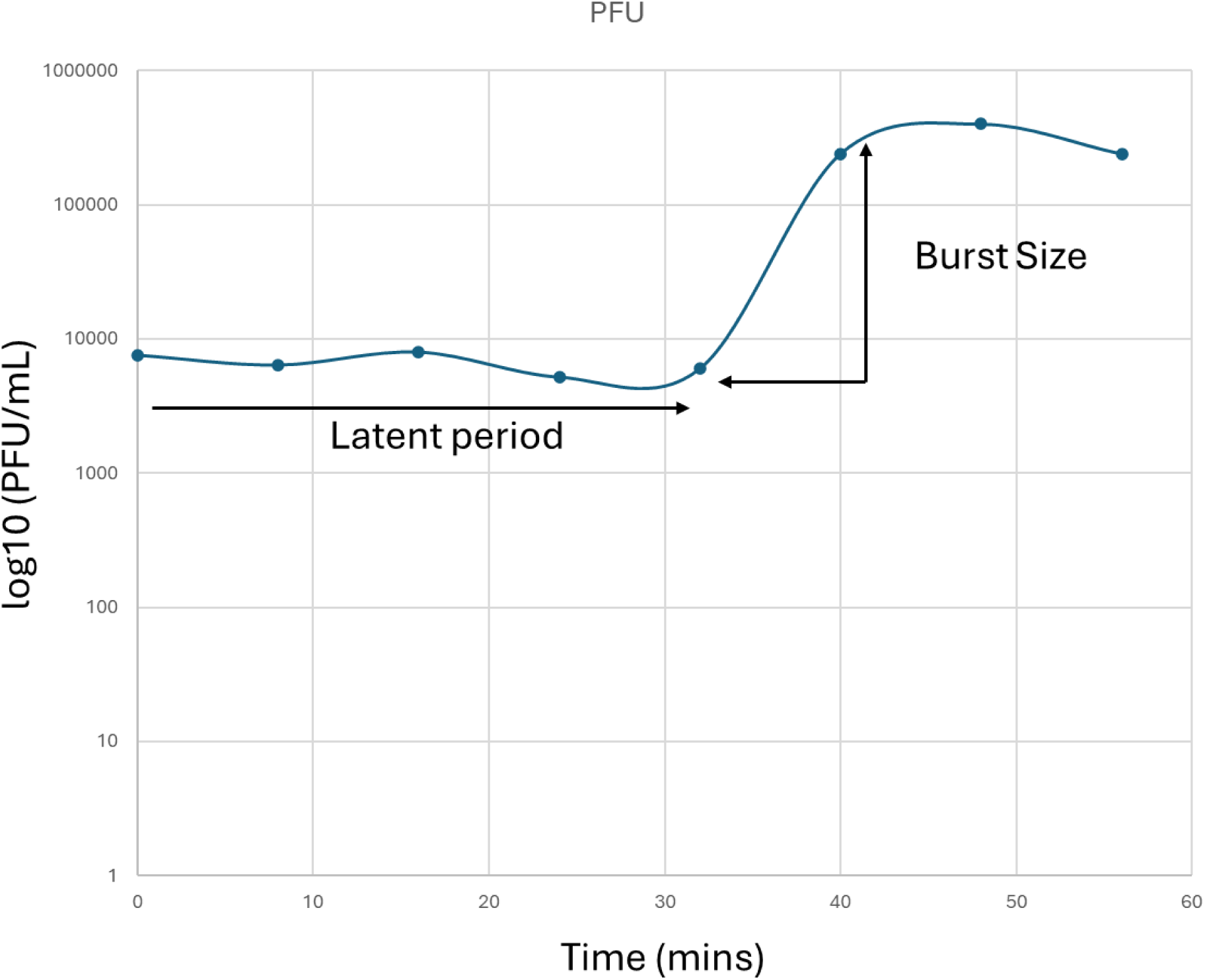
Calculation of the burst size and latent period/time to lysis to the cells by plotting the log of PFU against time.

The burst size for phage P278 was calculated at 46 and the time to first burst was observed to be ∼37 mins as seen in Figure S6.

## 3. Removal of static detections

We used a Density-Based Spatial Clustering of Applications with Noise (DBSCAN) algorithm to cluster detections that were spatially stable across time series so these incorrect detections could be excluded from the analysis. We set a threshold of 20% of detections over the entire time series found at the same location to classify a detection as static. They likely corresponded to false detections as cells are expected to be slightly motile and/or undergo Brownian motion. These detections were removed from the final cell counts.

**Figure S8:**
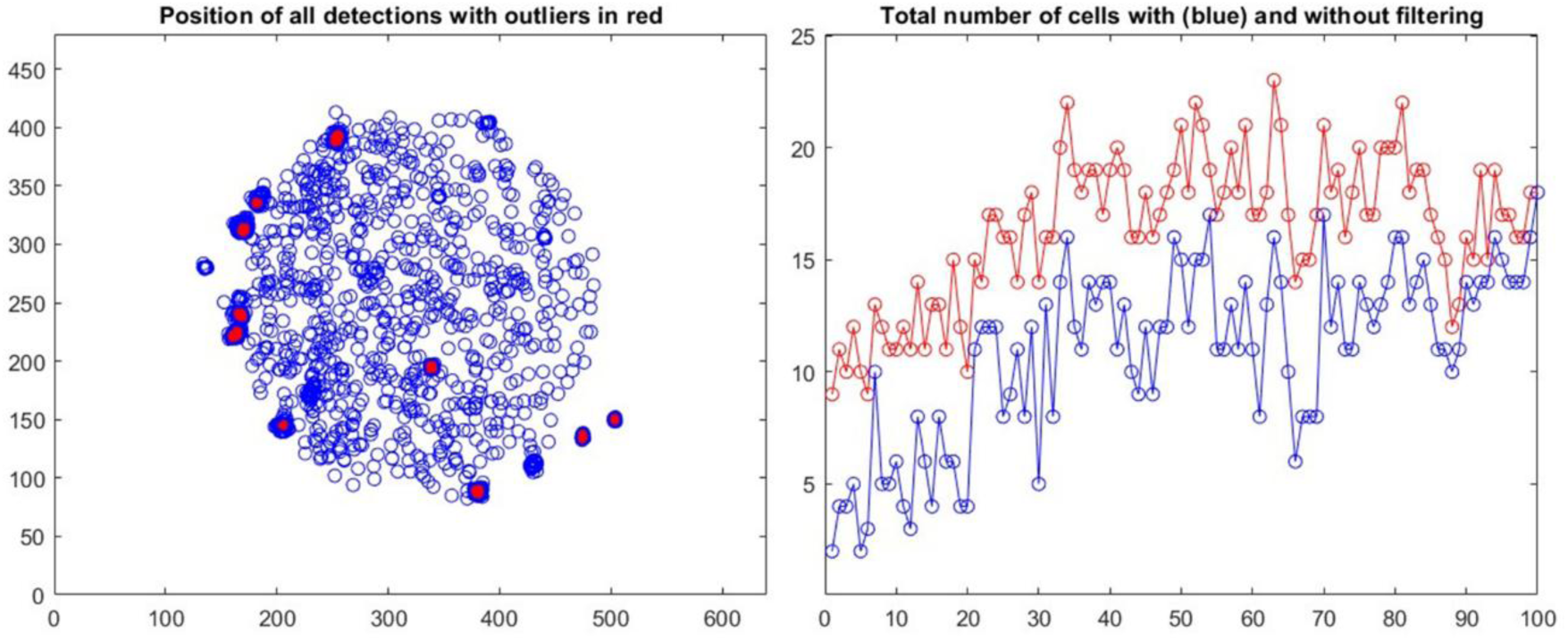
Removal of static and recurring false detections **A.** Detections remaining close to the same location are detected with a clustering algorithm. **B.** Example difference in counts with and without the static detection removal.

## 4. Time-lapse cell counts from individual droplets for PA-SA co-cultures

**Figure S9:**
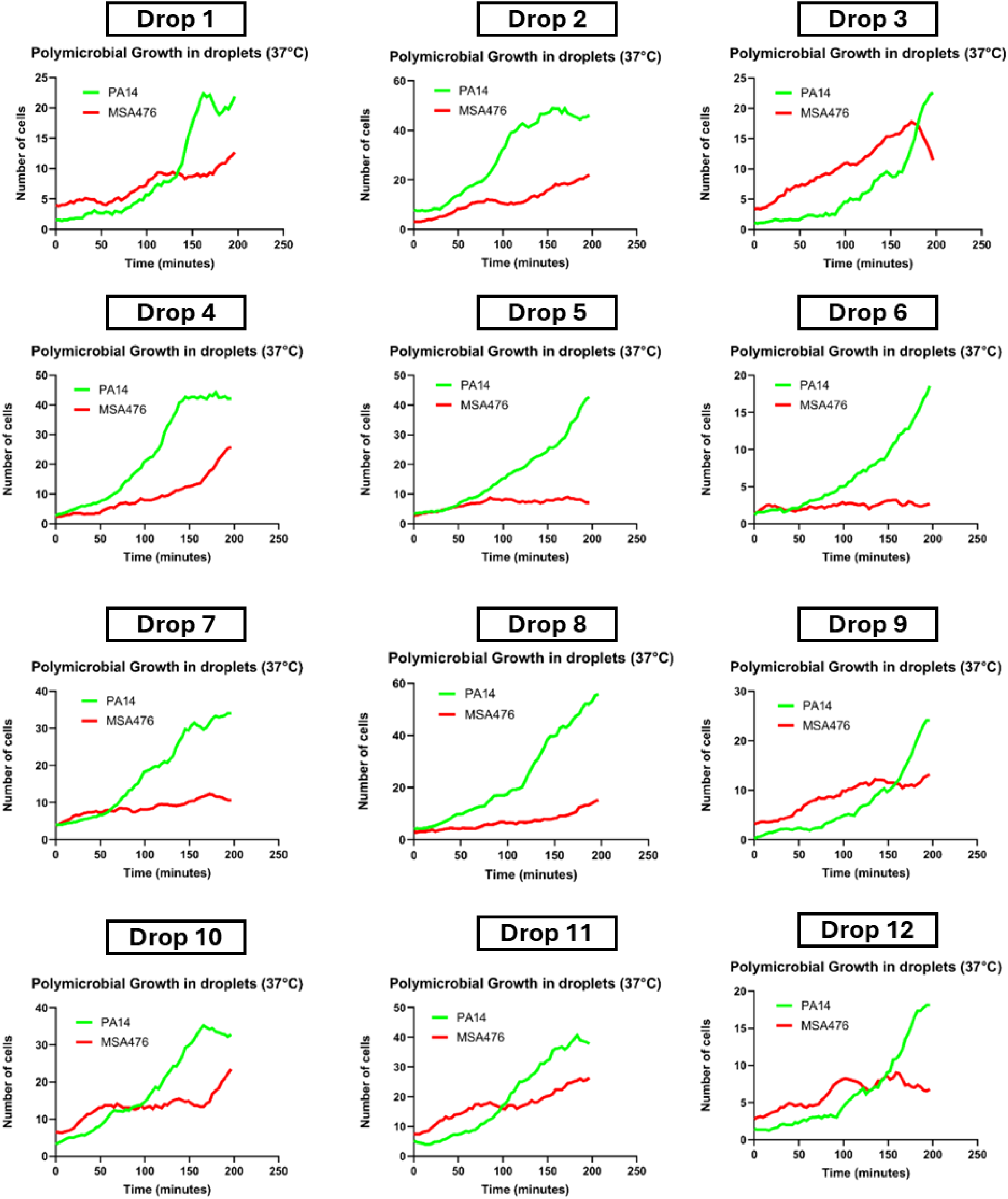
PA-SA co-culture growing in LB droplets showing the moving average of individual number of PA14 and MSSA cells observed over time in 12 droplets.

## 5. Time-lapse cell counts from individual droplets for bacteria-phage interactions

**Figure S10:**
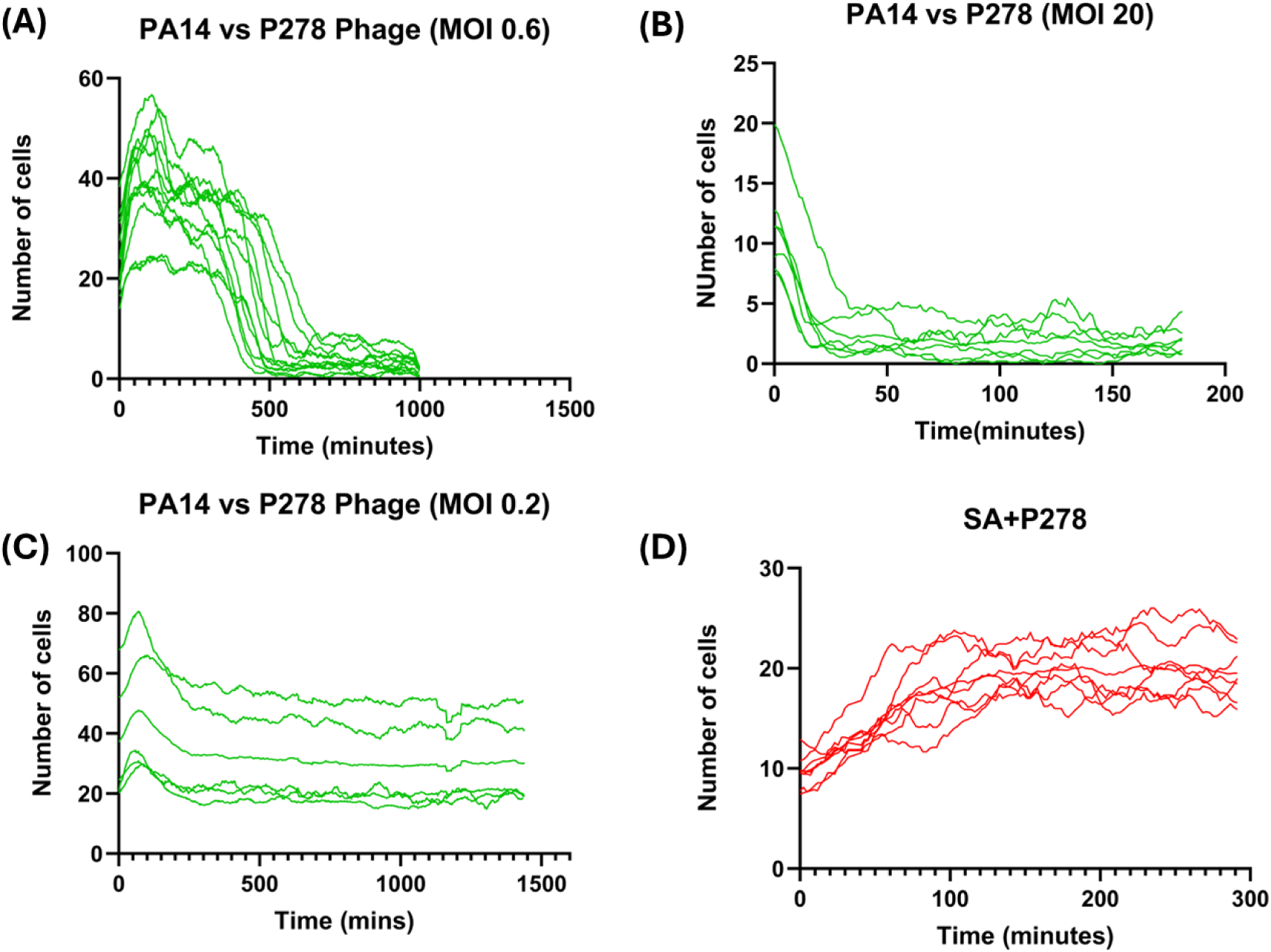
(A)-(C) Interaction of PA with P278 at different phage concentrations with varying cell numbers in droplet (D) Interaction of SA with phage P278 equivalent to an MOI of 0.2.

## 6. Time-lapse cell counts from individual droplets for two species and phage P278 interactions

**Figure S11:**
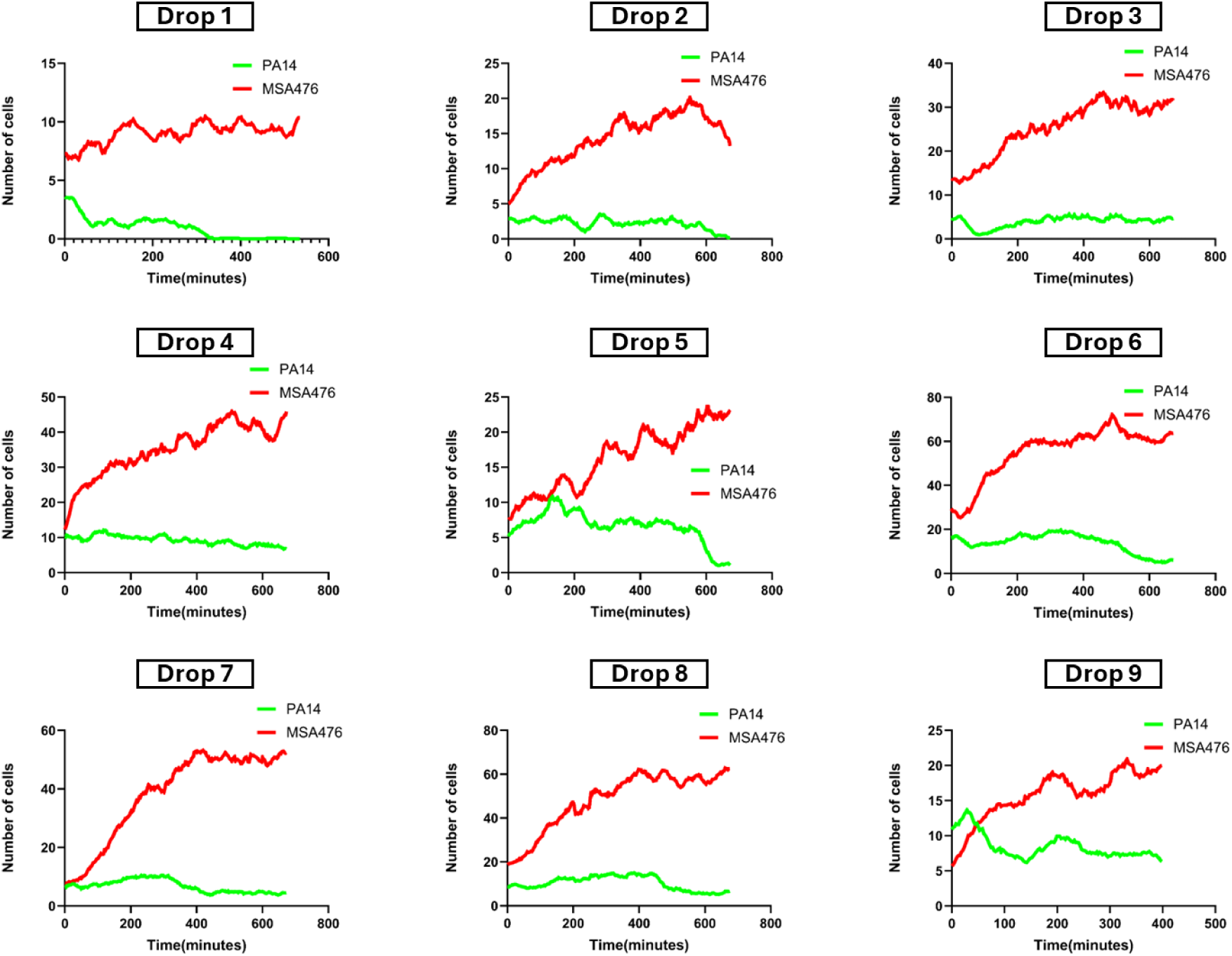
PA-SA co-culture with P278 phage in LB droplets showing the moving average of individual number of PA14 and MSSA cells observed over time in 9 droplets at MOI2.5.

**Figure S12:**
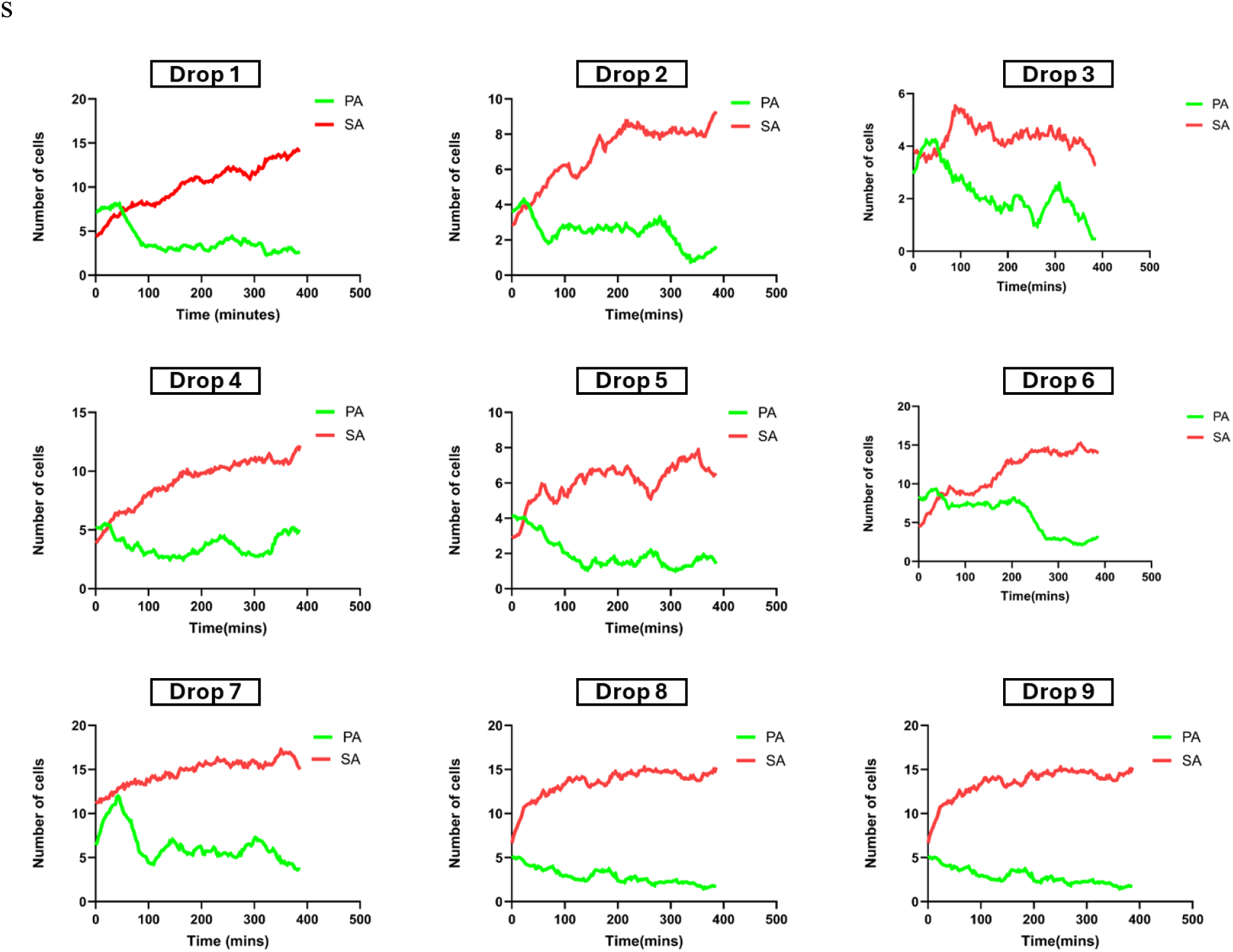
PA-SA co-culture with P278 phage in LB droplets showing the moving average of individual number of PA14 and MSSA cells observed over time in 9 droplets at MOI5.

